## Supporting Information for "Population genomics and molecular epidemiology of wheat powdery mildew in Europe"

Jigisha et al.

S3 Appendix. Recent evolution, population genetics and molecular epidemiology of *AvrPm17* 39

**References** **41**

### Tables

**S1 Table. Mating type frequency in the *Europe+_2022_2023* dataset**

Frequency of the two mating types MAT-1-2-1 and MAT-1-1-3 in the Bgt samples collected in 2022-2023 (See [1] for a characterization of the mating types). Mean freq. is the average frequency of the MAT-1-1-3 mating type in the population. The upper and lower 95% confidence intervals (UCI and LCI) and $p$-value were calculated with one-sample t-tests. The test was performed for populations that had more than 5 individuals.

| **Population** | **MAT-1-2-1** | **MAT-1-1-3** | **Mean freq** | **95% LCI** | **95% UCI** | ***p*-value** |
|---|---|---|---|---|---|---|
| N_EUR | 78 | 98 | 0.557 | 0.483 | 0.631 | 0.1321 |
| S_EUR1 | 6 | 6 | 0.5 | 0.168 | 0.832 | 1 |
| S_EUR2 | 30 | 29 | 0.491 | 0.36 | 0.623 | 0.898 |
| ME | 4 | 1 | 0.2 | - | - | - |
| TUR | 0 | 3 | 1 | - | - | - |
| **Total** | 118 | 137 | 0.537 | 0.476 | 0.599 | 0.235 |

**S2 Table. Demographic inference**

Results for the demographic analysis. The first two rows report the results for the datasets N_EUR2 and E_EUR2 assuming no misidentification of ancestral alleles. The last row reports the results for E_EUR2 allowing for misidentification of ancestral alleles (see **Methods**). The estimated parameters are $g$ (growth rate), $\alpha$ (parameter of the Beta coalescent), $e$ (probability of misidentification of the derived allele), and $\kappa$ (ratio between the mutation rates of transitions and transversions). The likelihood ratio (L. ratio) was calculated between the likelihood of the best model with $\alpha < 2$ (multiple merger) and the best model with $\alpha = 2$ (Kingman coalescent).

| **Dataset** | $g$ | $\alpha$ | $e$ | $\kappa$ | **L. ratio** |
|---|---|---|---|---|---|
| N_EUR_2 | 0 | 1.7 | - | - | 3605.9 |
| E_EUR_2 | 0.25 | 1.68 | - | - | 7257.6 |
| E_EUR_2 | 0.25 | 1.68 | 0.01 | 0.21 | 5388.7 |

**S3 Table. Mantel tests for isolation by geographic, wind and climatic distances**
The reported *p*-values are based on 999 permutations of the Mantel test. Pairwise genetic distances were computed as the number of SNPs between individuals scaled by the total number of loci compared. Geographic distance was calculated as the great circle distance between pairs of sampling locations, in kilometers. Wind distance was calculated as the mean estimated time of diffusion between a pair of sampling locations (in wind hours). Climatic distance was calculated as the Euclidean distance between pairs of samples based on the first seven principal components of the PCA on 12 climatic variables (see **Methods**).

| Dataset | Test | Mantel correlation | *p*-value |
|---|---|---|---|
| Europe+_2022_2023 | Isolation by geographic distance | 0.472 | 0.001 |
| Europe+_2022_2023 | Isolation by wind distance | 0.547 | 0.001 |
| Europe+_2022_2023 | Isolation by climatic distance | 0.33 | 0.001 |
| N_EUR_2022_2023 | Isolation by geographic distance | 0.085 | 0.011 |
| N_EUR_2022_2023 | Isolation by wind distance | 0.129 | 0.01 |
| N_EUR_2022_2023 | Isolation by climatic distance | -0.049 | 0.849 |
| S_EUR2_2022_2023 | Isolation by geographic distance | 0.389 | 0.001 |
| S_EUR2_2022_2023 | Isolation by wind distance | 0.259 | 0.001 |
| S_EUR2_2022_2023 | Isolation by climatic distance | 0.342 | 0.001 |

**S4 Table. Correlation between geographic, wind and climatic distances**
The reported *p*-values are based on 999 permutations of the Mantel Test. All pairwise distance values in column "x,y" are the same as described in **S3 Table**.

| Dataset | x, y | Mantel correlation | *p*-value |
|---|---|---|---|
| Europe+_2022_2023 | Geography, Wind | 0.788 | 0.001 |
| Europe+_2022_2023 | Geography, Climate | 0.6 | 0.001 |
| N_EUR_2022_2023 | Geography, Wind | 0.65 | 0.001 |
| N_EUR_2022_2023 | Geography, Climate | 0.731 | 0.001 |
| S_EUR2_2022_2023 | Geography, Wind | 0.829 | 0.001 |
| S_EUR2_2022_2023 | Geography, Climate | 0.557 | 0.001 |

**S5 Table. Logistic regression to test for effects of geographic, wind and climatic distances** Response variable 'Diff pop' was 0 if a pair of individuals belonged to the same population and 1 if not. Geographic, wind and climate distances for all pairs of individuals were scaled and centered. The analysis was performed for all isolates of the *Europe+_2022_2023* dataset that belonged to either the N_EUR population or to S_EUR2 (n= 235, number of pairs of individuals = 27495). Nagelkerke $R^2$ values are reported for each model. The first three rows show results of the simple regression models where each variable was tested separately. The 'FULL' model is the multiple regression model with all three variables.

| **Model** | **Intercept** | **Coefficient** | ***p*-value** | $\mathbf{R^2}$ |
|---|---|---|---|---|
| Diff pop ∼ geographic distance | -0.599 | 1.2 | <0.0001 | 0.3 |
| Diff pop ∼ wind distance | -0.6389 | 1.879 | <0.0001 | 0.497 |
| Diff pop ∼ climate distance | -0.6272 | 1.176 | <0.0001 | 0.276 |
| Diff pop ∼ geographic distance + wind distance + climate distance (*FULL*) | -0.7355 | - | - | 0.533 |
| *FULL*: geographic distance | - | -0.388 | <0.0001 | - |
| *FULL*: wind distance | - | 1.878 | <0.0001 | - |
| *FULL*: climate distance | - | 0.754 | <0.0001 | - |

**S6 Table. Test for isolation by distance along the east-west and north-south axes** The reported *p*-values are based on 999 permutations of the Mantel Test. For details on the calculation of pairwise geographic distances along the two axes, see **Methods**.

| **Dataset** | **Axis** | **Mantel correlation** | ***p*-value** |
|---|---|---|---|
| N_EUR_2022_2023 | East-West | 0.028 | 0.203 |
| N_EUR_2022_2023 | North-South | 0.125 | 0.007 |
| S_EUR2_2022_2023 | East-West | 0.394 | 0.001 |
| S_EUR2_2022_2023 | North-South | 0.119 | 0.018 |

**S7 Table. Variance partitioning using ANOVA following RDA**

This analysis was performed for the *Europe+_recent* dataset. 'Genotypes' is the binary genotype matrix of filtered SNPs. Climate represents the 12 shortlisted climate variables. Wind stands for the first three wind coordinates (see **Methods**). Geography is the sampling location (latitude, longitude) of the isolate and country is the country of sampling. Inertia can be interpreted as variance.

| **Model** | **Inertia** | ***p*-value** | **Proportion of explainable variance** | **Proportion of total variance ($R^2$)** | **Adjusted $R^2$** |
|---|---|---|---|---|---|
| Full $genotypes \sim climate + wind + geography + country$ | 3939.4 | <0.001 | 1 | 0.202 | 0.11 |
| Climate $genotypes \sim climate\vert(wind + geography + country)$ | 848.15 | <0.001 | 0.215 | 0.043 | 0.015 |
| Wind $genotypes \sim wind\vert(climate + geography + country)$ | 225.54 | <0.001 | 0.057 | 0.012 | 0.005 |
| Geography $genotypes \sim geography\vert(wind + climate + country)$ | 158.59 | <0.001 | 0.04 | 0.008 | 0.004 |
| Country $genotypes \sim country\vert(wind + climate + geography)$ | 1265.16 | <0.001 | 0.321 | 0.065 | 0.015 |
| Confounded | 1441.96 | - | 0.366 | 0.074 | - |
| Total unexplained | 15587.32 | - | - | 0.798 | - |
| Total inertia | 19526.72 | - | - | 1 | - |

**S8 Table. Variance partitioning using ANOVA following RDA including host species**
This analysis was performed for a subset of the *Europe+_2022_2023* dataset containing isolates that had been sampled from infected fields with known host ploidy (i.e. hexaploid or tetraploid wheat; n=131). 'Genotypes' is the binary genotype matrix of filtered SNPs. Climate represents the 12 shortlisted climate variables. Wind stands for the first three wind coordinates (see **Methods**). Geography is the sampling location (latitude, longitude) of the isolate and country is the country of sampling. Host reflects the wheat ploidy - 6n or 4n. Inertia can be interpreted as variance.

| Model | Inertia | *p*-value | Proportion of explainable variance | Proportion of total variance ($R^2$) | Adjusted $R^2$ |
|---|---|---|---|---|---|
| Full<br>$genotypes \sim climate + wind + geography + country + host$ | 14101.08 | <0.001 | 1 | 0.316 | 0.084 |
| Climate<br>$genotypes \sim climate \vert (wind + geography + country + host)$ | 3690.72 | 0.779 | 0.262 | 0.083 | -0.002 |
| Wind<br>$genotypes \sim wind \vert (climate + geography + country + host)$ | 961.18 | 0.35 | 0.068 | 0.022 | 0.001 |
| Geography<br>$genotypes \sim geography \vert (wind + climate + country + host)$ | 646.3 | 0.337 | 0.046 | 0.015 | 0.001 |
| Country<br>$genotypes \sim country \vert (wind + climate + geography + host)$ | 4819.47 | 0.13 | 0.342 | 0.108 | 0.003 |
| Host<br>$genotypes \sim host \vert (wind + climate + geography + country)$ | 428.02 | <0.001 | 0.03 | 0.01 | 0.003 |
| Confounded | 3555.39 | | 0.252 | 0.08 | |
| Total unexplained | 30462.84 | | | 0.684 | |
| Total inertia | 44563.92 | | | 1 | |

**S9 Table. AvrPm17 protein variants**

Protein variants are defined based on the amino acid (aa) sequence of the mature protein (excluding the first 25 aa coding for the signal peptide). For variant nomenclature we follow [2]. There are six polymorphic aa in the mature protein, for each of these positions we report which aa is present in which variant. Additionally, we report the number of isolates carrying each variant in the *Europe+* dataset.

| **AvrPm17 variant** | **31** | **53** | **55** | **61** | **80** | **103** | **N. of isolates in *Bgt_Europe+*** |
|---|---|---|---|---|---|---|---|
| Variant A | Y | A | E | G | R | S | 70 |
| Variant B | - | V | - | - | S | - | 125 |
| Variant C | - | V | - | - | - | - | 246 |
| Variant F | - | V | - | - | - | R | 2 |
| Variant H | H | V | - | - | - | - | 12 |
| Variant I | H | V | - | - | S | - | 1 |
| Variant J | - | - | - | - | S | - | 1 |
| Variant K | - | - | R | A | - | - | 1 |
| Variant L | - | - | R | A | S | - | 1 |

**S10 Table. Association between AvrPm17 protein variants and sampling period**

For this analysis we used 74 samples collected from Switzerland, France and the UK. Samples were divided into two periods based on their year of collection: samples collected between 1980 and 2001 (Before 2002), and samples collected in 2022-2023. The variant counts represent the number of isolates that carried the respective variant (in at least one copy). We tested whether the frequency of a variant was associated with a specific period using the Fisher exact test. The odd ratio represents the over- or under-representation of a variant in the earlier period compared to the recent period.

| **AvrPm17 variant** | **Before 2002** | **2022-2023** | **Odd ratio** | ***p*-value** |
|---|---|---|---|---|
| Variant A | 6 (30%) | 6 (9.5%) | 3.99 | 0.034 |
| Variant B | 2 (10%) | 9 (14.3%) | 0.67 | 0.475 |
| Variant C | 11 (55%) | 45 (71.4%) | 0.49 | 0.138 |
| Variant H | 0 | 2 (3.2%) | 0 | 0.574 |
| Other variants | 1 | 1 | - | - |
| Total | 20 | 63 | - | - |

**S11 Table. Association between AvrPm17 protein variants and relatedness clusters**
Total clustered isolates are all isolates belonging to one of the nine largest clusters. Total non-clustered isolates are all isolates that are not identical-by-descent over the AvrPm17 locus with any other sample (singletons). We counted how many isolates carried each variant, independently on the number of copies. The odd ratio is the over- or under-representation of a variant in clustered strains compared to non-clustered strains. The $p$-values were computed with one-sided Fisher exact tests.

| AvrPm17 variant | Clustered isolates | Non-clustered isolates | Odd ratio | $p$-value |
|---|---|---|---|---|
| Variant A | 35 | 3 | 2.62 | 0.075 |
| Variant B | 75 | 16 | 0.97 | 0.524 |
| Variant C | 168 | 22 | 2.32 | 0.002 |
| Variant H | 11 | 1 | 2.35 | 0.355 |
| Two different variants | 14 | 21 | 0.10 | 1.7e-09 |
| Missing data | 3 | 1 | | |
| Total | 306 | 64 | | |

### Figures

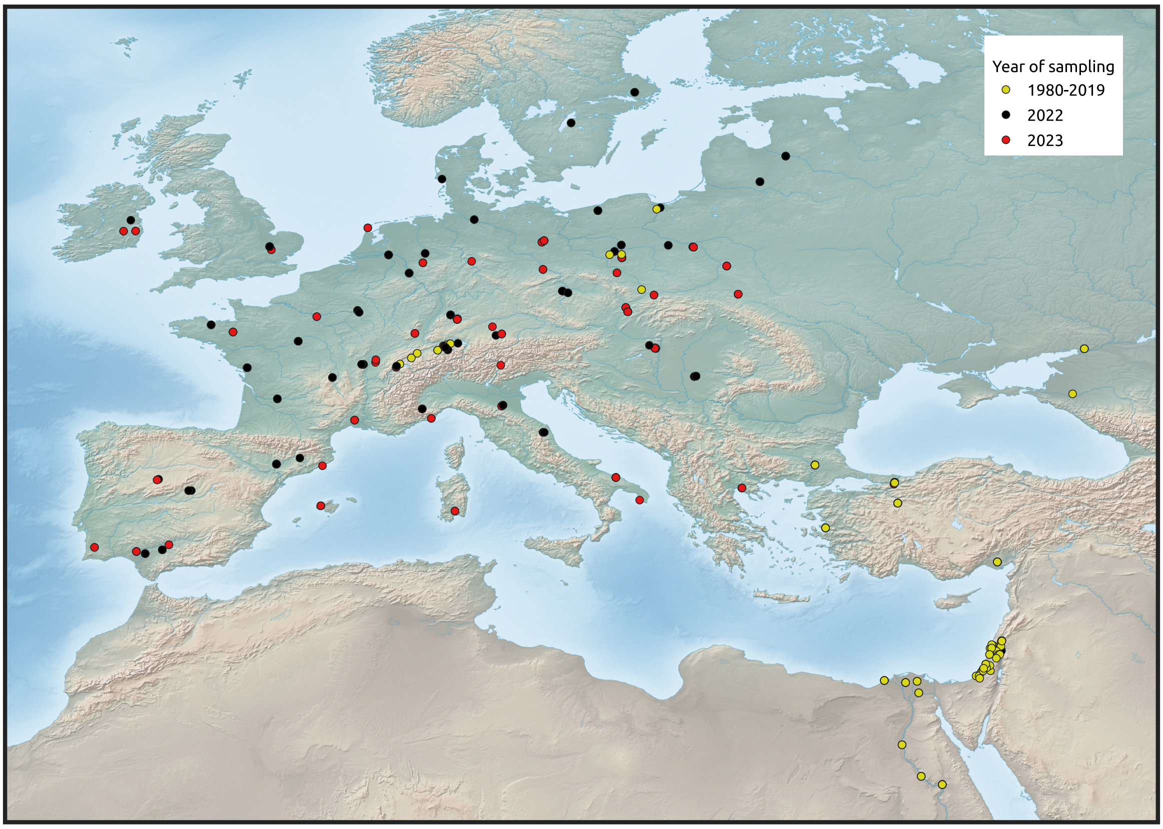

**S1 Fig. Sampling locations of Bgt isolates in the *Europe+* dataset**
The black and red circles correspond to sampling locations of the isolates collected in 2022 and 2023. In yellow are the sampling locations of the isolates collected between 1980-2019 whose whole-genome sequences were publicly available. Samples in the *Europe+* dataset with no available geo-spatial data are not shown on the map.

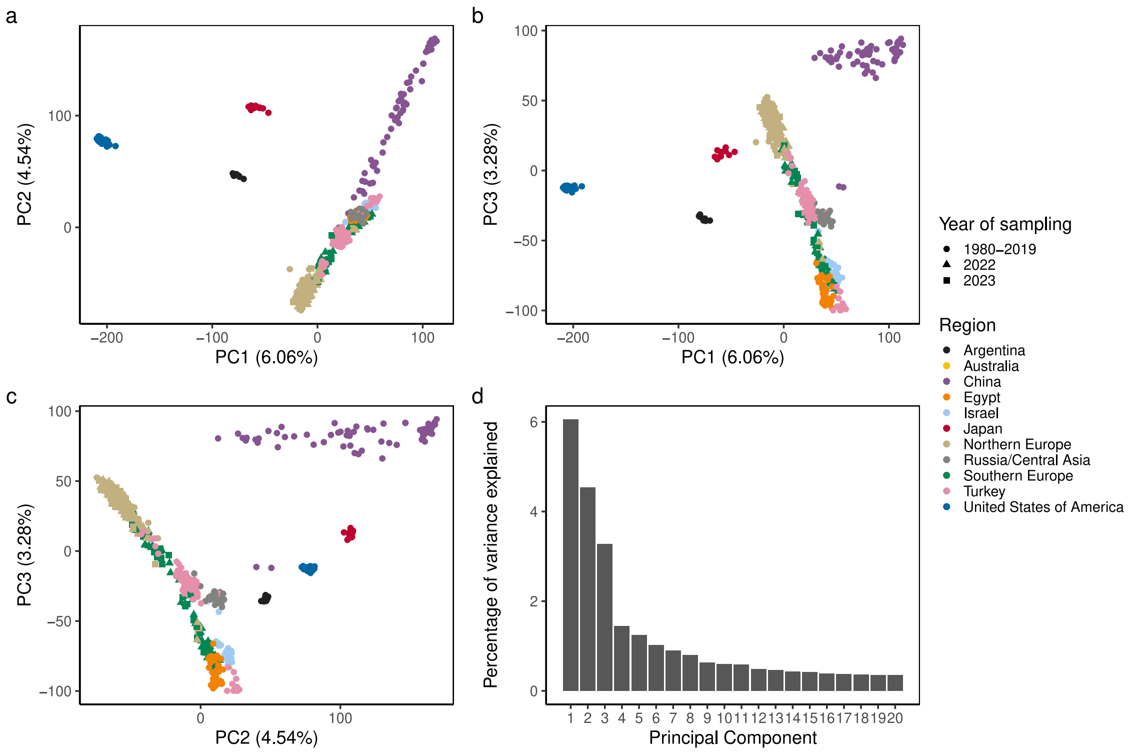

**S2 Fig. Principal component analysis (PCA) of the *World* dataset**
**(a)** PC1 vs. PC2 **(b)** PC1 vs. PC3 **(c)** PC2 vs. PC3. Each point represents an individual and the colours indicate the respective regions of sampling. The shape of the symbols represents the year the isolate was sampled in. **(d)** Percentage of variance explained by the first 20 principal components.

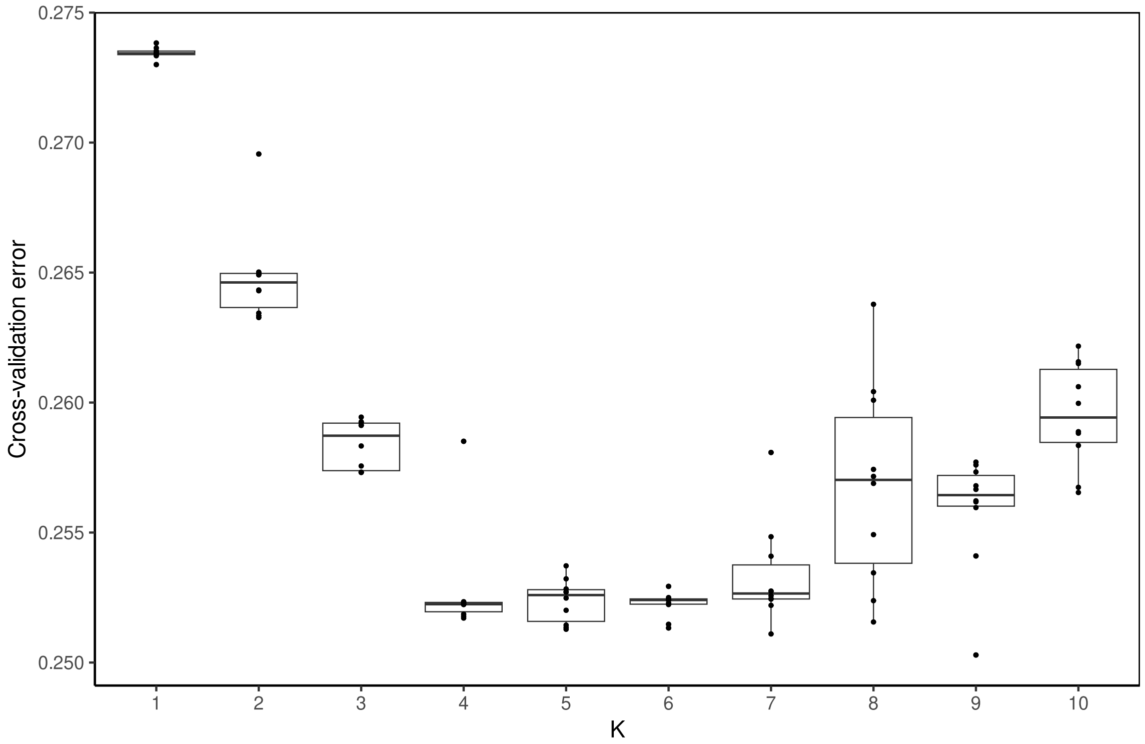

**S3 Fig. ADMIXTURE cross-validation error for the *World* dataset**
The ADMIXTURE model was run for 10 replicates for each value of K from 1 to 10. The cross-validation errors from all runs are plotted, with each dot representing one run. Boxes show the inter-quartile range of CV-error for each K, with the horizontal line representing the median.

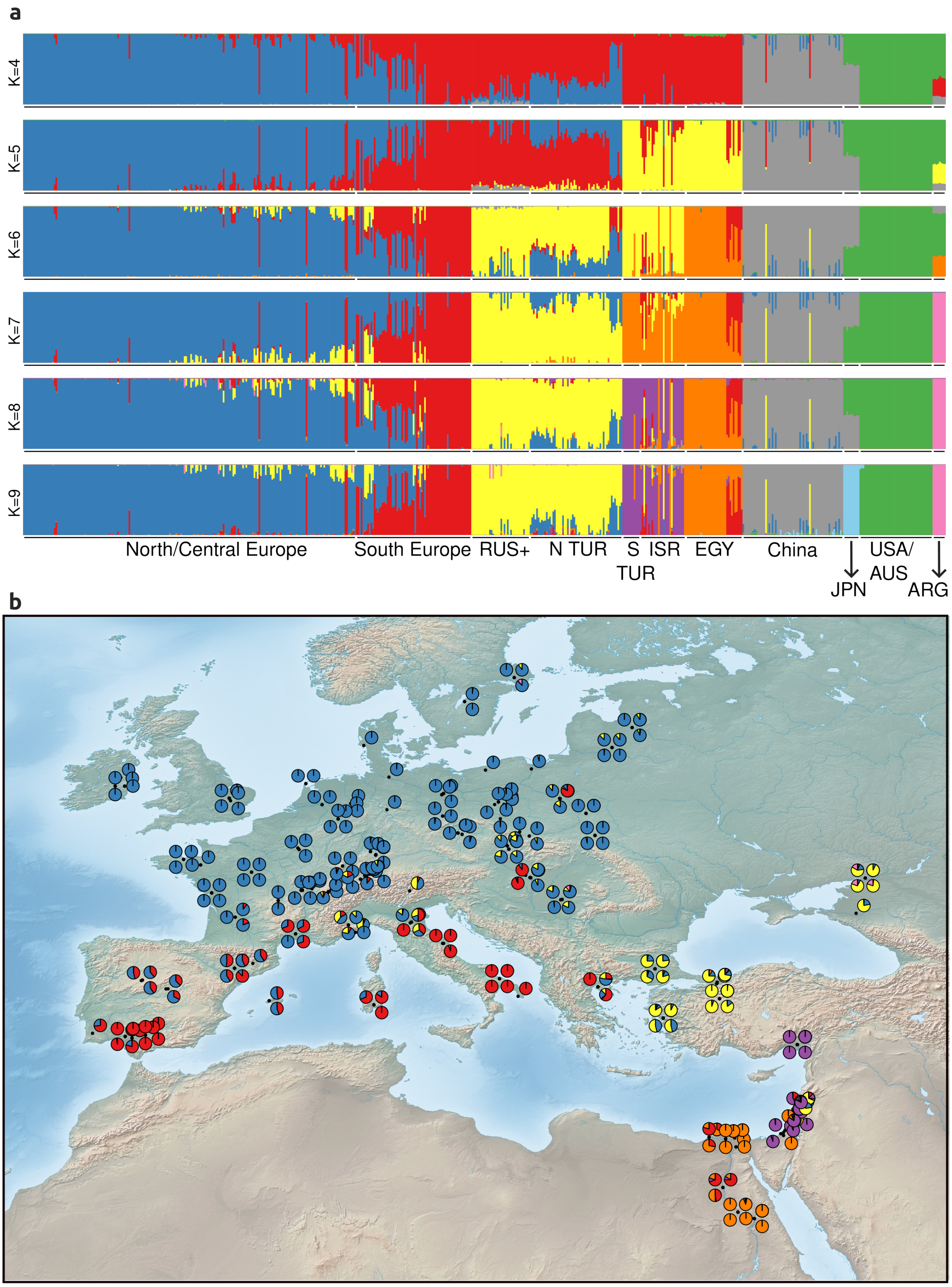

**S4 Fig. Individual ancestry proportions inferred by ADMIXTURE**
(**a**) Ancestry proportions of the *World* dataset obtained from the replicate with the least cross-validation error for a given value of K are plotted for K=4-9. Each bar represents an individual and each unique colour represents a different ancestry. RUS- Russia, N TUR- Northern Turkey, S TUR- Southern Turkey, ISR- Israel, EGY- Egypt, JPN- Japan, USA/AUS- USA/Australia, ARG- Argentina. (**b**) Map showing ancestry proportions of individuals belonging to the *Europe+* dataset. The ancestry proportion values are taken from (**a**), panel K=9. The black dots denote the sampling locations, and each pie chart represents the proportion of ancestries for a given individual. Samples in the *Europe+* dataset with no available geo-spatial data are not shown on the map. For graphical reasons, at most four individuals are shown for any location.

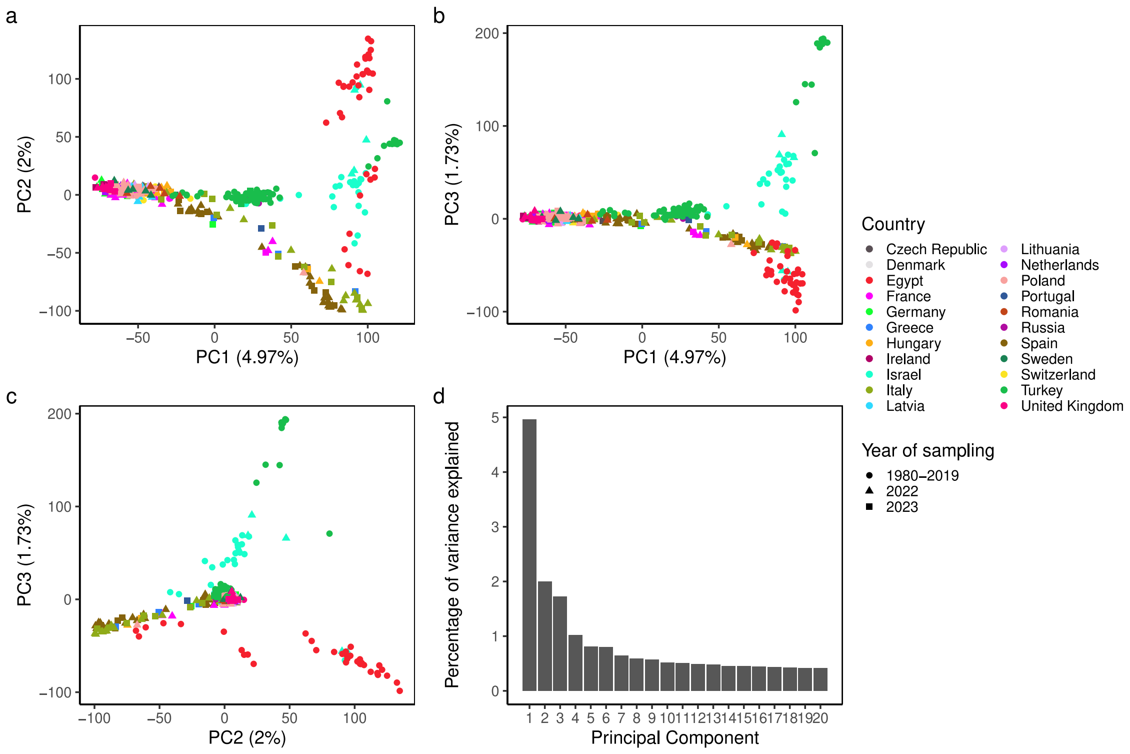

**S5 Fig. Principal component analysis (PCA) of the *Europe+* dataset**
(**a**) PC1 vs. PC2 (**b**) PC1 vs. PC3 (**c**) PC2 vs. PC3. Each point represents an individual coloured by the country it was sampled from. The shape of the symbols represents the year the isolate was sampled in. (**d**) Percentage of variance explained by the first 20 principal components.

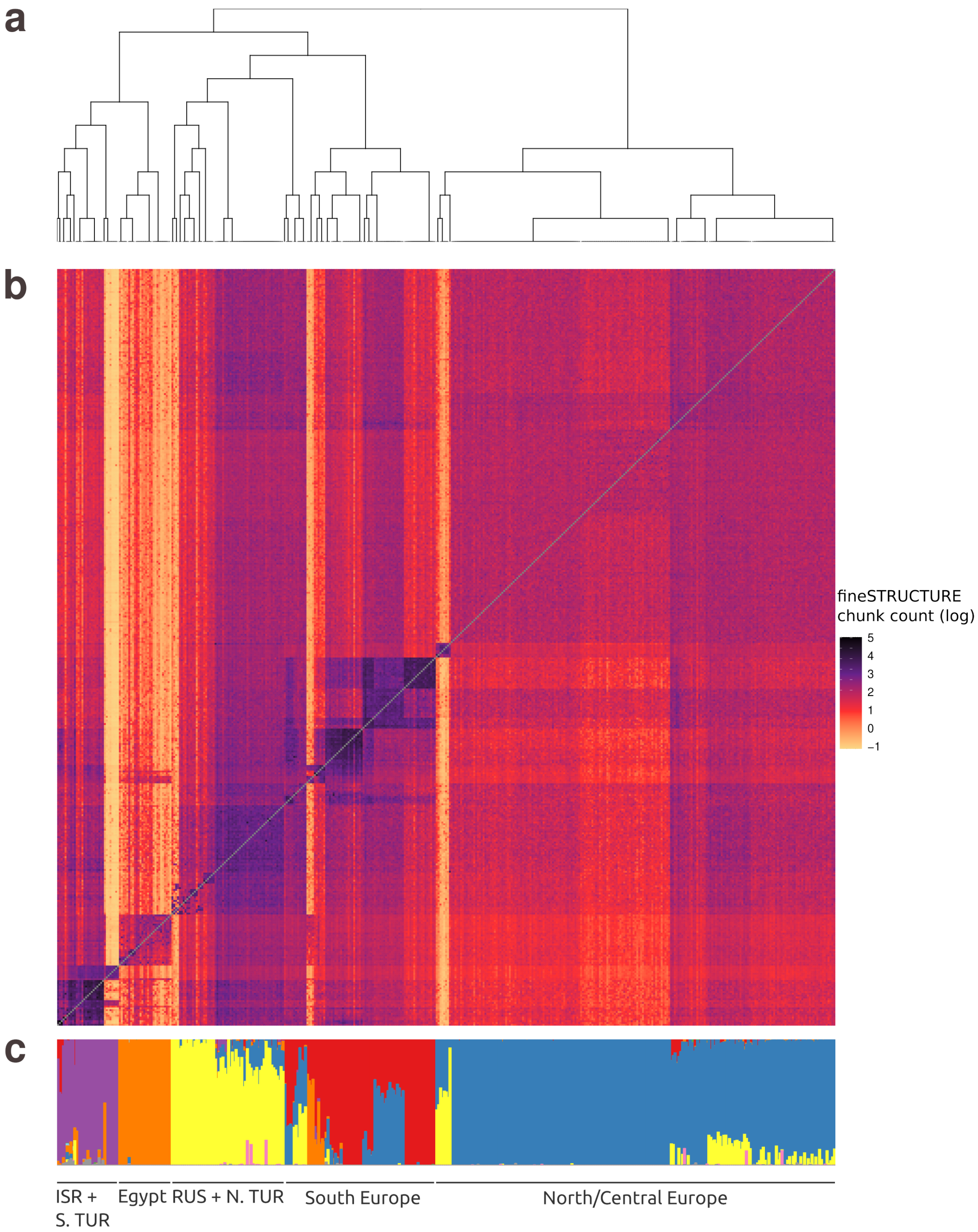

**S6 Fig. fineSTRUCTURE analysis of the *Europe+* dataset**
(**a**) Dendrogram and (**b**) coancestry matrix computed by fineSTRUCTURE. Colours in the matrix represent chunk count (in log scale) which is the number of genomic segments donated by isolates in rows to isolates in columns. Darker colours indicate higher relatedness. (**c**) ADMIXTURE barplot representing individual ancestry proportions, same as panel K=9 in **S4a Fig**. The order of the samples in (**a**), (**b**) and (**c**) is the same so that one column represents one sample throughout the figure. ISR + S. TUR- Israel + South Turkey, RUS + N. TUR- Russia + North Turkey.

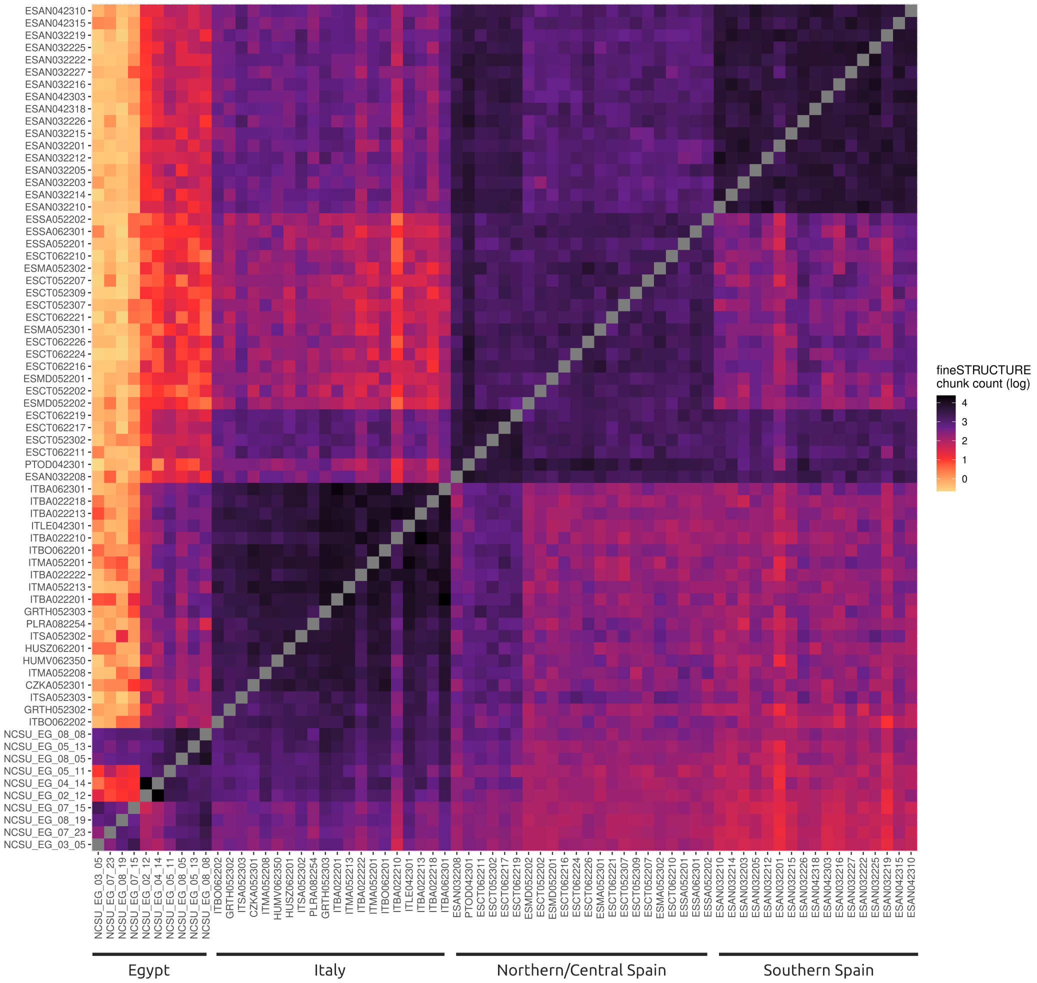

**S7 Fig. Coancestry matrix for Southern Europe**

Subset of the fineSTRUCTURE coancestry matrix of the *Europe+* dataset to show isolates belonging to the Southern European population (S_EUR2). Chunk count is the number of genomic segments donated by isolates in rows to isolates in columns (in log scale). Darker colours indicate higher relatedness.

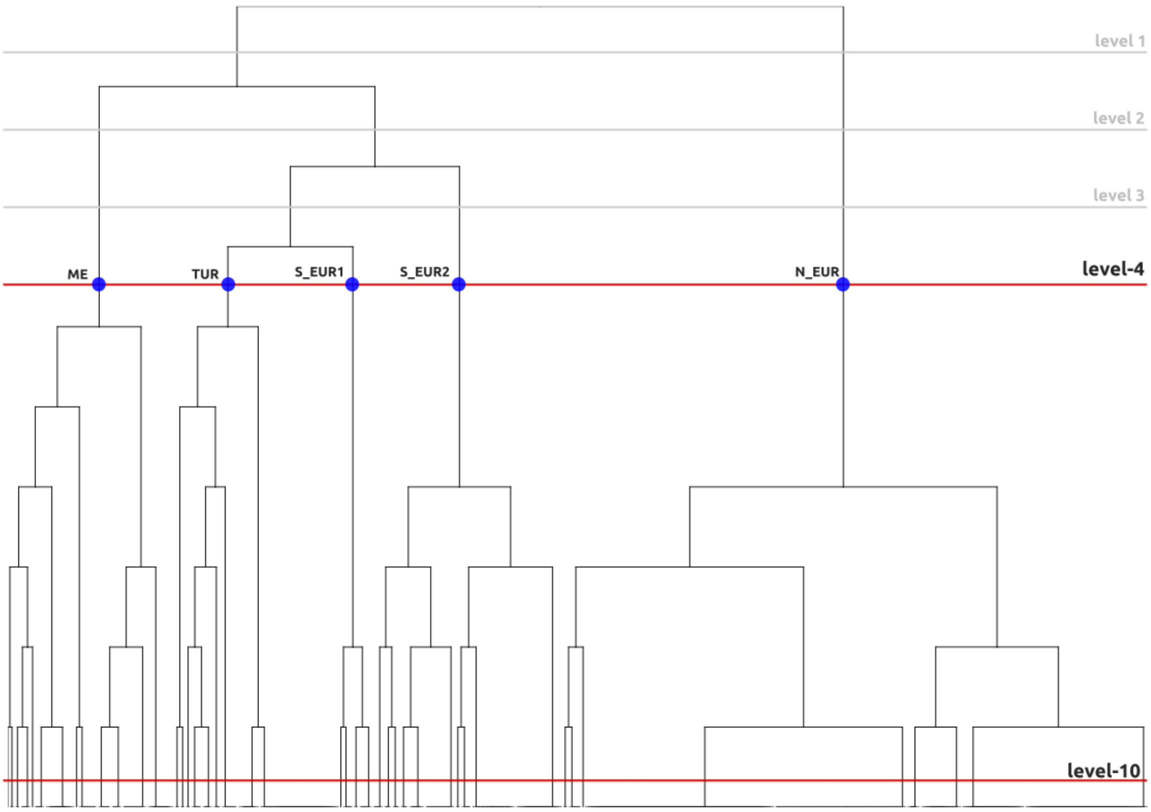

**S8 Fig. Population classification based on the fineSTRUCTURE dendrogram**
Population dendrogram, inferred by fineSTRUCTURE (same as in **Fig 1b** and **S6b Fig**). Slicing the dendrogram at different levels groups the *Europe+* dataset into different numbers of populations. At each level, individuals belonging to the same population share more ancestry with each other than they do with individuals from other populations. The ‘level-4’ grouping divides the data into 5 populations – ME, TUR, S_EUR1, S_EUR2 and N_EUR – and was used for most population-level analyses. At the finest level, ’level-10’, there are 45 populations (from left to right: ISR1, ISR2, ISR3, ISR4, ISR5, ISR+TUR, ISR6, TUR1, TUR2, EG1, EG2, EG_ISR, EG3, TUR3, TUR4, TUR5, TUR6, TUR7, TUR8, TUR9, TUR10, TUR11, S_EUR1, S_EUR2, S_EUR3, S_EUR4, EG4, EG5, EG6, IT+_1, IT+_2, IT, SPAIN_N1, SPAIN_N3, SPAIN_N2, SPAIN_S, TUR12, TUR13, TUR14, N_EUR1, N_EUR2, N_EUR_old+, N_EUR_old, E_EUR1, E_EUR2). ’Level-10’ was used for the demographic analysis (**S2 Table and S12 Fig**), and to study spatio-temporal patterns of genetic variation (**S16-17 Fig**).

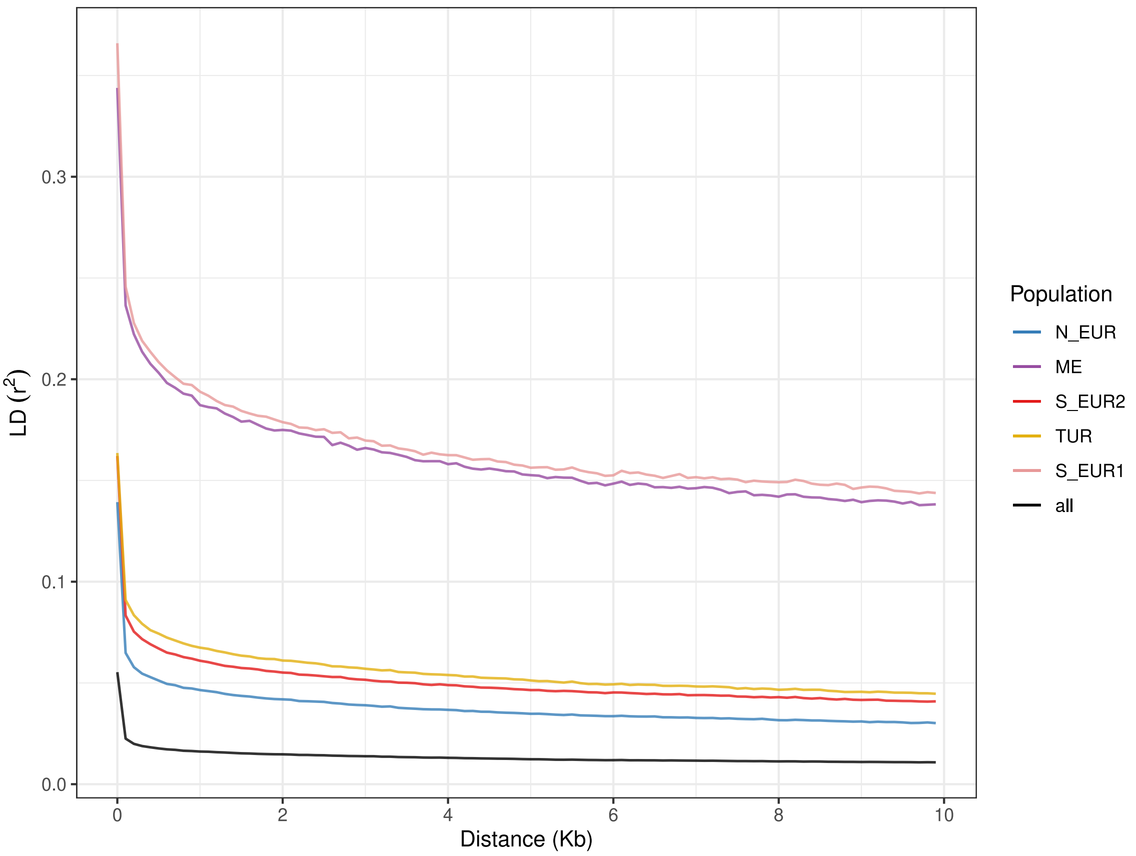

**S9 Fig. Linkage disequilibrium decay**
Linkage disequilibrium decay (measured as $r^2$) within the five populations (N_EUR, S_EUR2, TUR, S_EUR1, and ME) and across the *Europe+_recent* dataset (all).

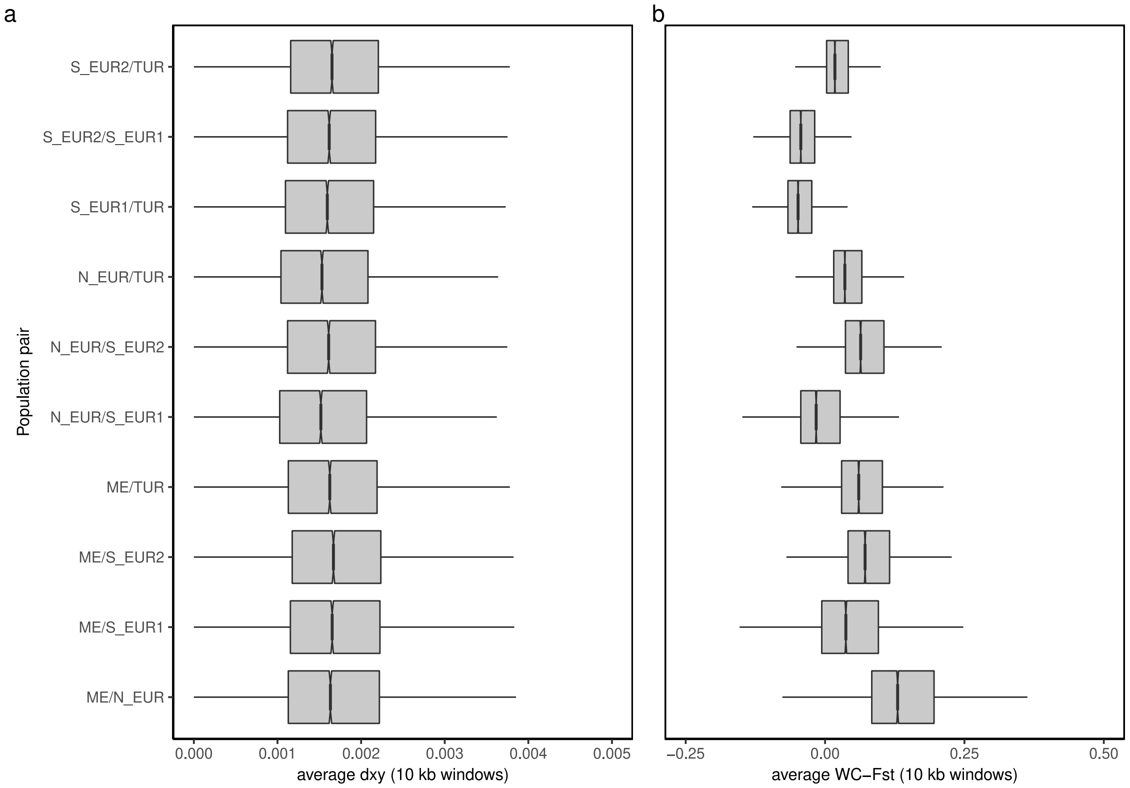

**S10 Fig. Summary statistics for pairwise population differentiation**
(**a**) Average per-site between population nucleotide divergence, $d_{xy}$, and (**b**) average per-SNP Weir and Cockerham's $F_{ST}$ calculated for all pairs of populations over the whole genome in windows of 10 kb.

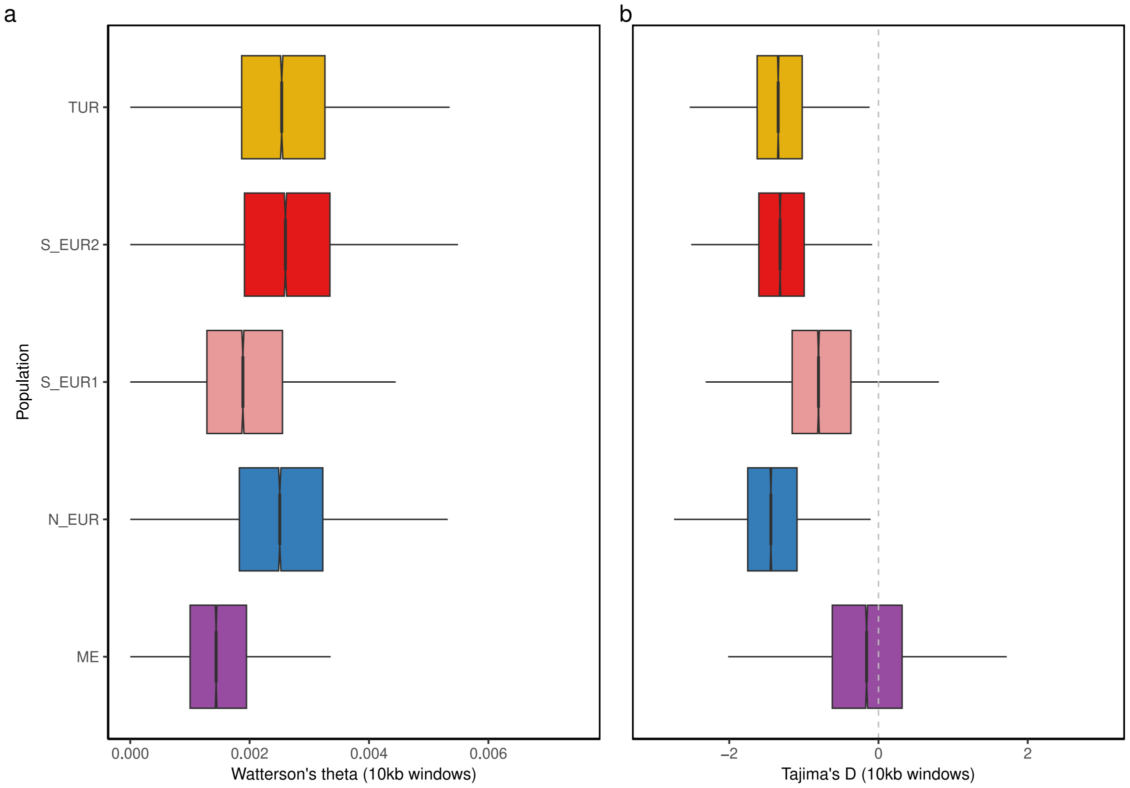

**S11 Fig. Genome-wide population summary statistics**
Population-level averages of (**a**) Watterson's theta and (**b**) Tajima's D calculated for the five populations (N_EUR, S_EUR2, TUR, S_EUR1, and ME) over the whole genome in windows of 10 kb.

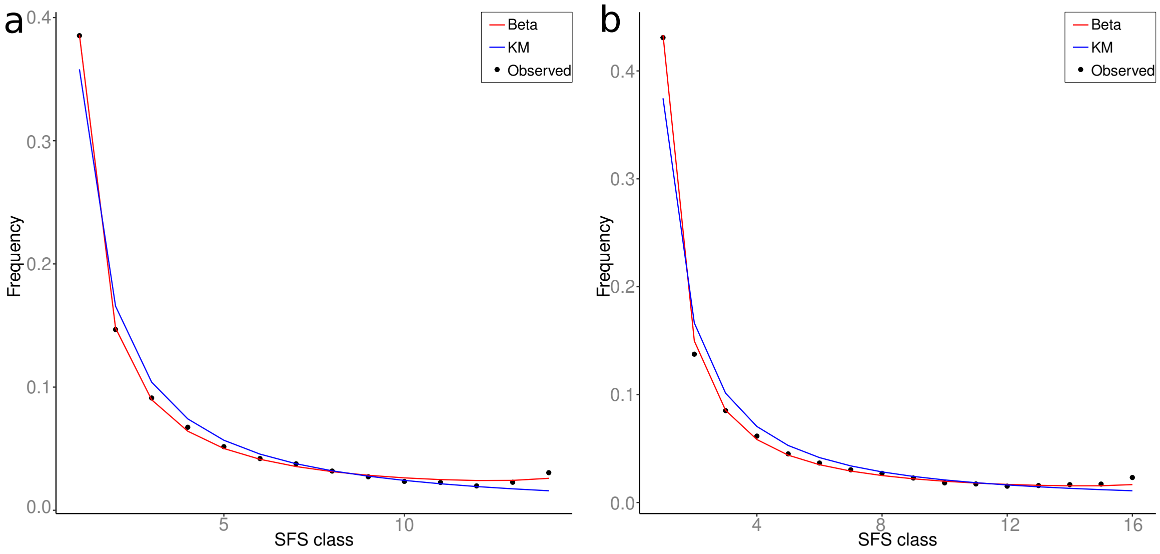

**S12 Fig. Site frequency spectra for N_EUR2 and E_EUR2 populations**
The observed site frequency spectra are depicted with black dots. The expected values for the best fitting multiple merger model and the best fitting Kingman model are shown with red and blue lines respectively. (**a**) Population N_EUR2 (fineSTRUCTURE level-10), and (**b**) population E_EUR2 (fineSTRUCTURE level-10).

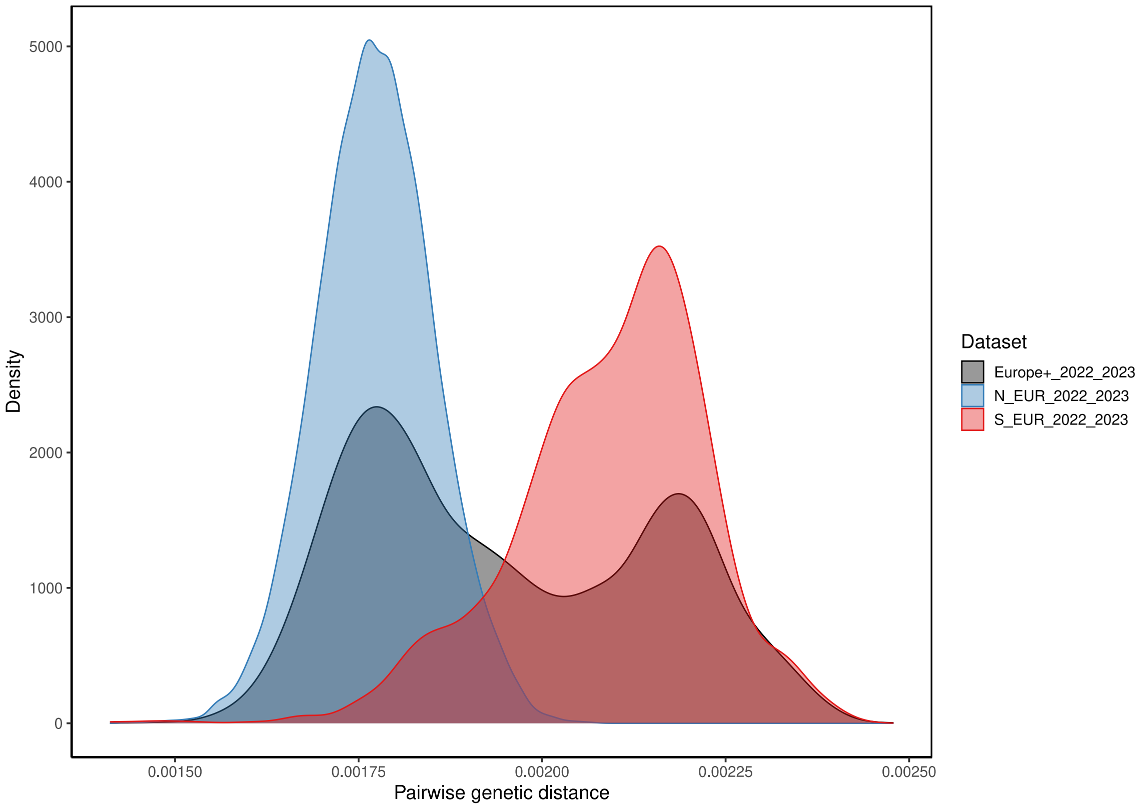

**S13 Fig. Distribution of pairwise genetic distances**
Kernel density estimates of pairwise genetic distances for the *Europe+_2022_2023* dataset (grey) and for samples of this dataset belonging to the populations N_EUR (blue) and S_EUR2 (red).

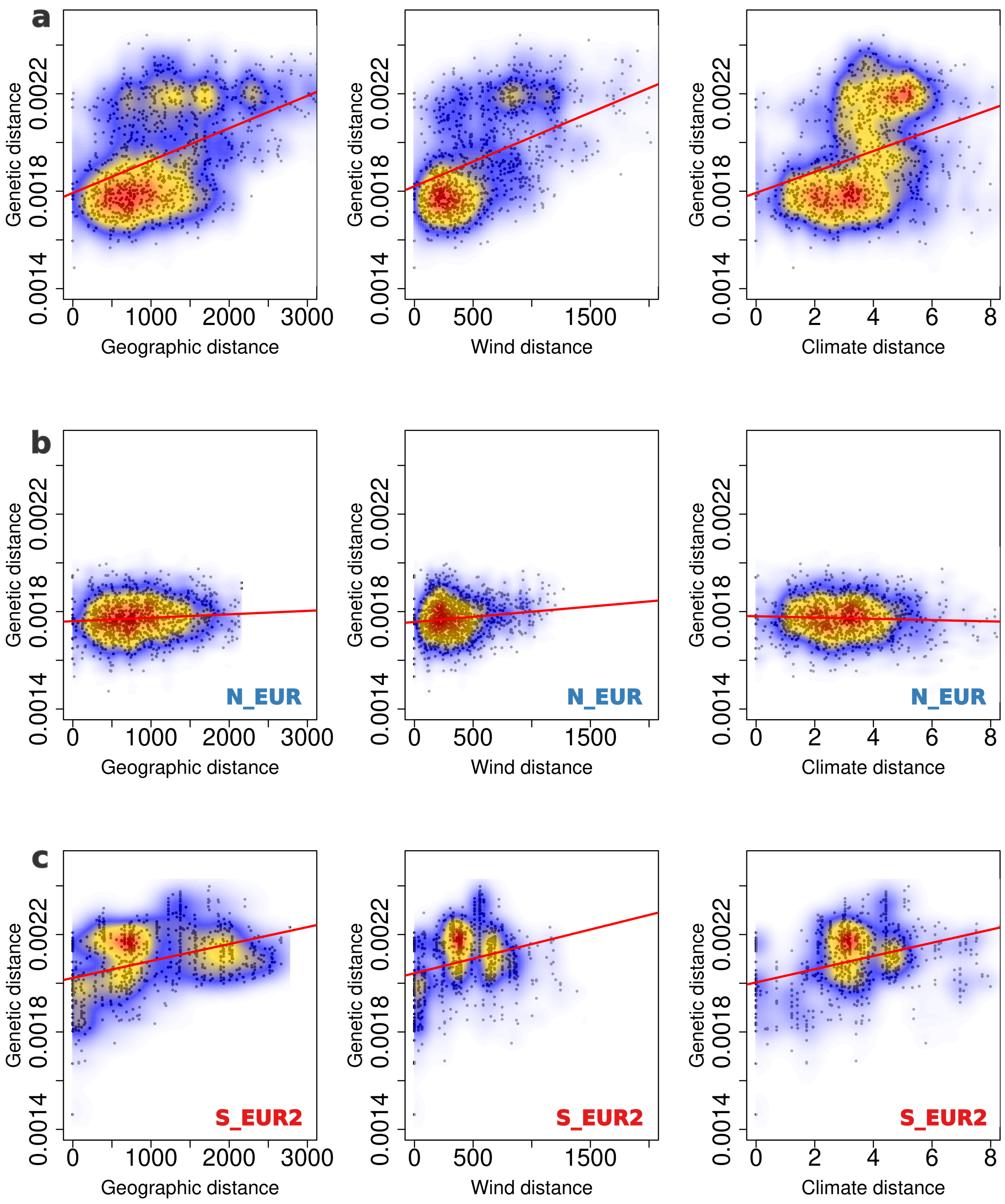

**S14 Fig. Isolation by geographic, wind and climatic distances**
Pairwise genetic distances plotted against pairwise geographic, wind and climatic distances in columns 1, 2 and 3 respectively for (**a**) *Europe+_2022_2023*, (**b**) isolates belonging to N_EUR in the *Europe+_2022_2023* dataset, and (**c**) isolates belonging to S_EUR2 in the *Europe+_2022_2023* dataset. Genetic distances were calculated as the number of SNPs between individuals scaled by the total number of loci compared. Geographic distance was measured as the great circle distance between sampling locations, in kilometers. Wind distances were estimated as the time of diffusion between pairs of sampling locations, in wind hours. Climatic distances were calculated as the euclidean distance between samples based on the first seven principal components of the PCA on the 12 shortlisted climatic variables (see **Methods**). The colours represent the density of the data points, with warmer colours corresponding to higher density. Only 1000 randomly sampled data points for each dataset are plotted in black. The corresponding Mantel test results are reported in **S3 Table**.

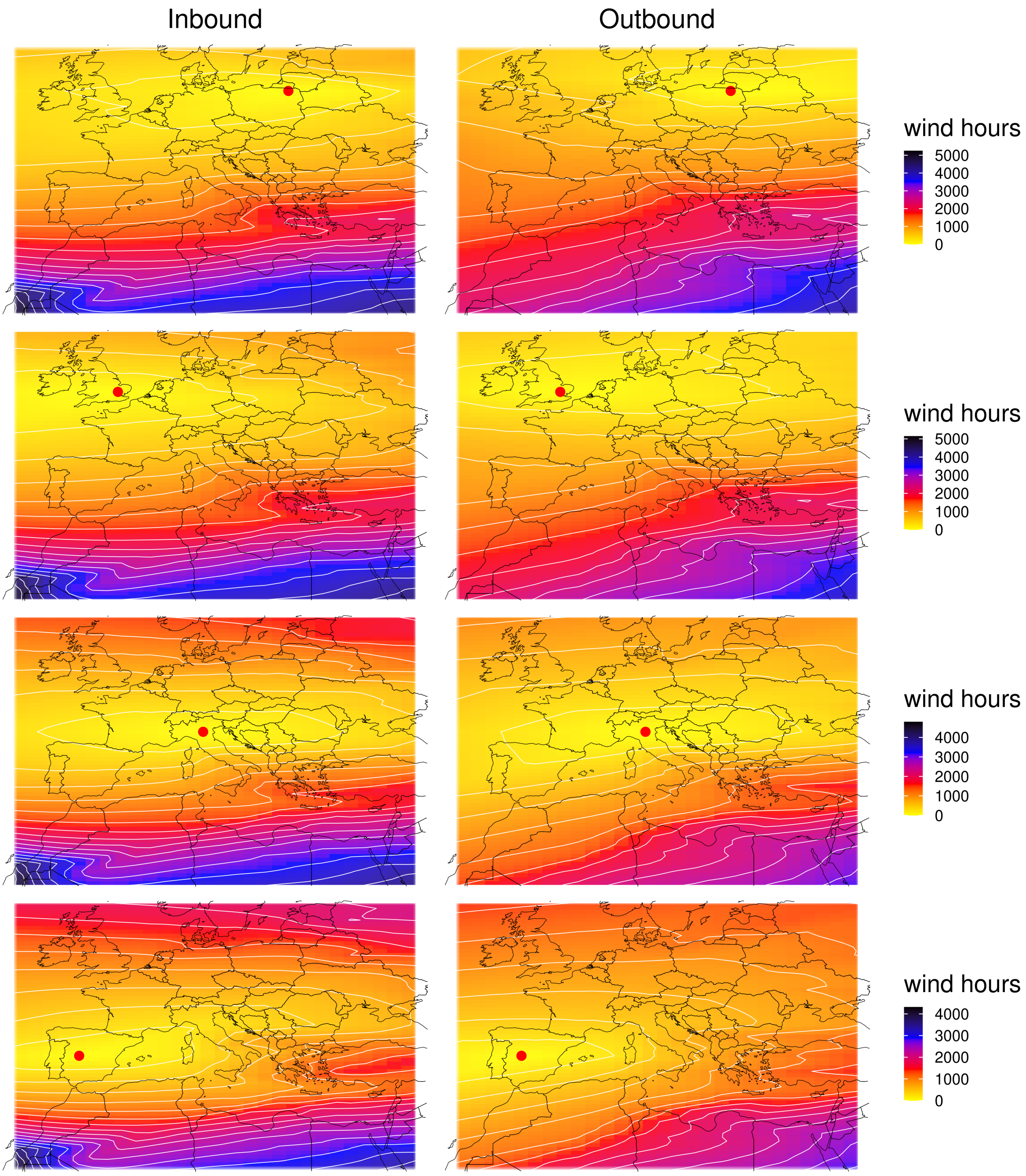

**S15 Fig. Wind connectivity surfaces for four selected locations**
Inbound and Outbound connectivity surfaces based on wind data from 2012 to 2021 for four focal locations (red circles). The darker the colour the longer is the average time needed to travel to, or from the focal locality.

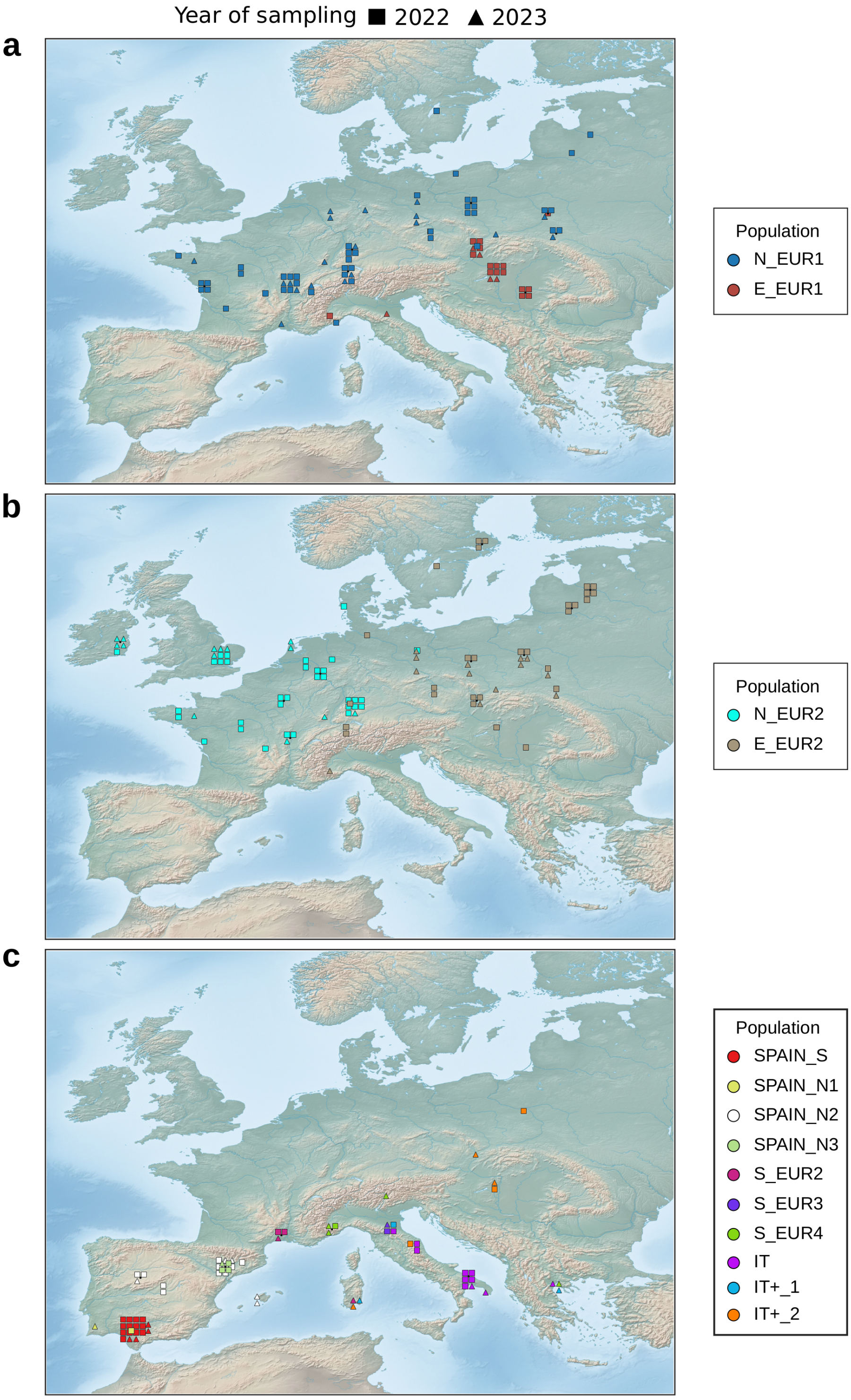

**S16 Fig. Temporal persistence of local populations from 2022 to 2023**
Examples of fineSTRUCTURE level-10 populations in northern Europe (**a** and **b**) and southern Europe (**c**) illustrating the similarity between individuals sampled in 2022 (squares) and 2023 (triangles) from nearby locations. Only samples collected in these two years are shown on the maps. Each colour represents a different population.

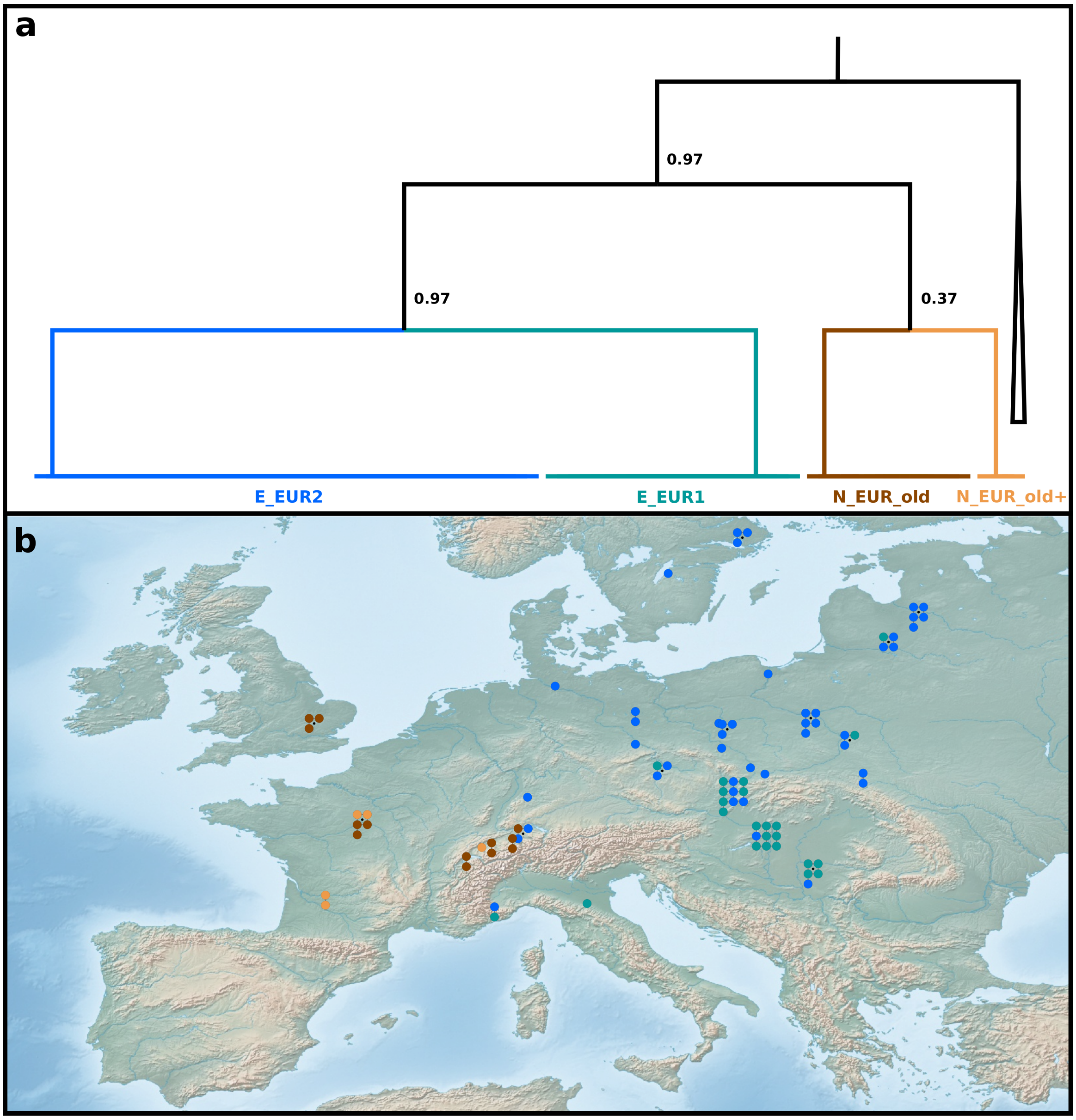

**S17 Fig. Geographic shift of Bgt populations over two decades**
(**a**) Subtree of the fineSTRUCTURE dendrogram with fineSTRUCTURE level-10 populations labelled. N_EUR_old is the only population composed entirely of older samples from the UK, France, and Switzerland (all collected before 2000, except one Swiss isolate sampled in 2007). N_EUR_old+ is composed of three older samples from Switzerland and France, plus two isolates collected in the south of France in 2022. E_EUR1 and E_EUR2 are composed of recent samples collected after 2017 (mostly in 2022 and 2023). The node labels represent fineSTRUCTURE merging probabilities for the descendant populations. (**b**) Geographic distribution of the isolates composing E_EUR2, E_EUR1, N_EUR_old, and N_EUR_old+. Colours as in panel **a**.

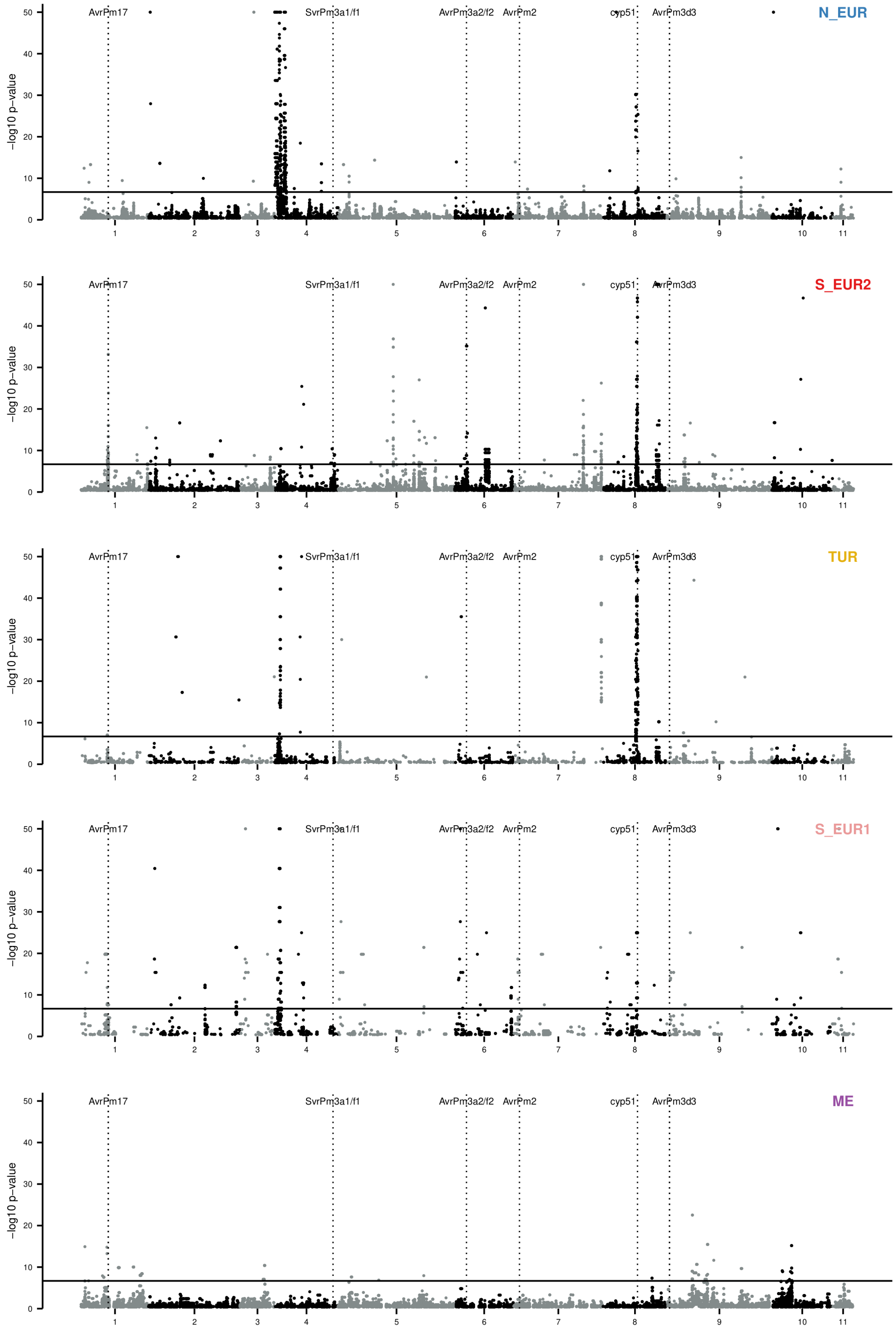

**S18 Fig. Genome-wide scans for signatures of recent selection**
Manhattan plot produced by the isoRelate analysis of the five populations (N_EUR, S_EUR2, TUR, S_EUR1, and ME). The y axis represents the significance of the excess of relatedness in a population. SNPs in regions with a significant excess of identity by descent pairs indicate loci under positive selection in the recent past ($\sim$ 25 sexual generations). The horizontal full lines show the Bonferroni corrected 0.05 threshold. Vertical dotted lines show the position of five known avirulence genes [2, 3, 4, 5] and the fungicide target cyp51. SNPs with -log $p$-values less than 0.4 are not shown.

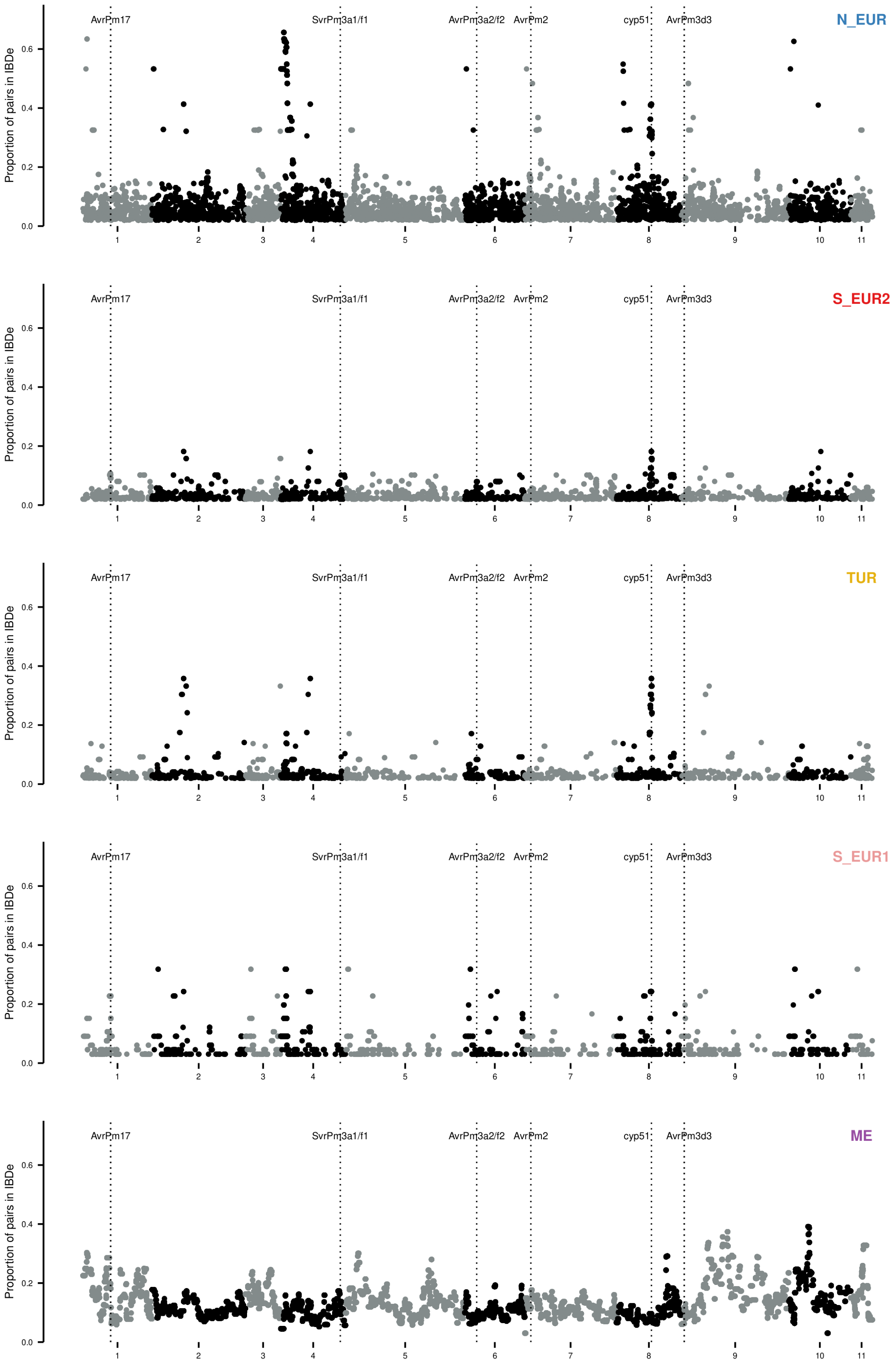

**S19 Fig. Genome-wide pairwise relatedness**
Manhattan plot for the isoRelate analysis of the five populations (N_EUR, S_EUR2, TUR, S_EUR1, and ME). The y axis represents the proportion of pairs of isolates identitical-by-descent (IBDe) within a population. SNPs with IBDe proportion less than 0.02 are not shown.

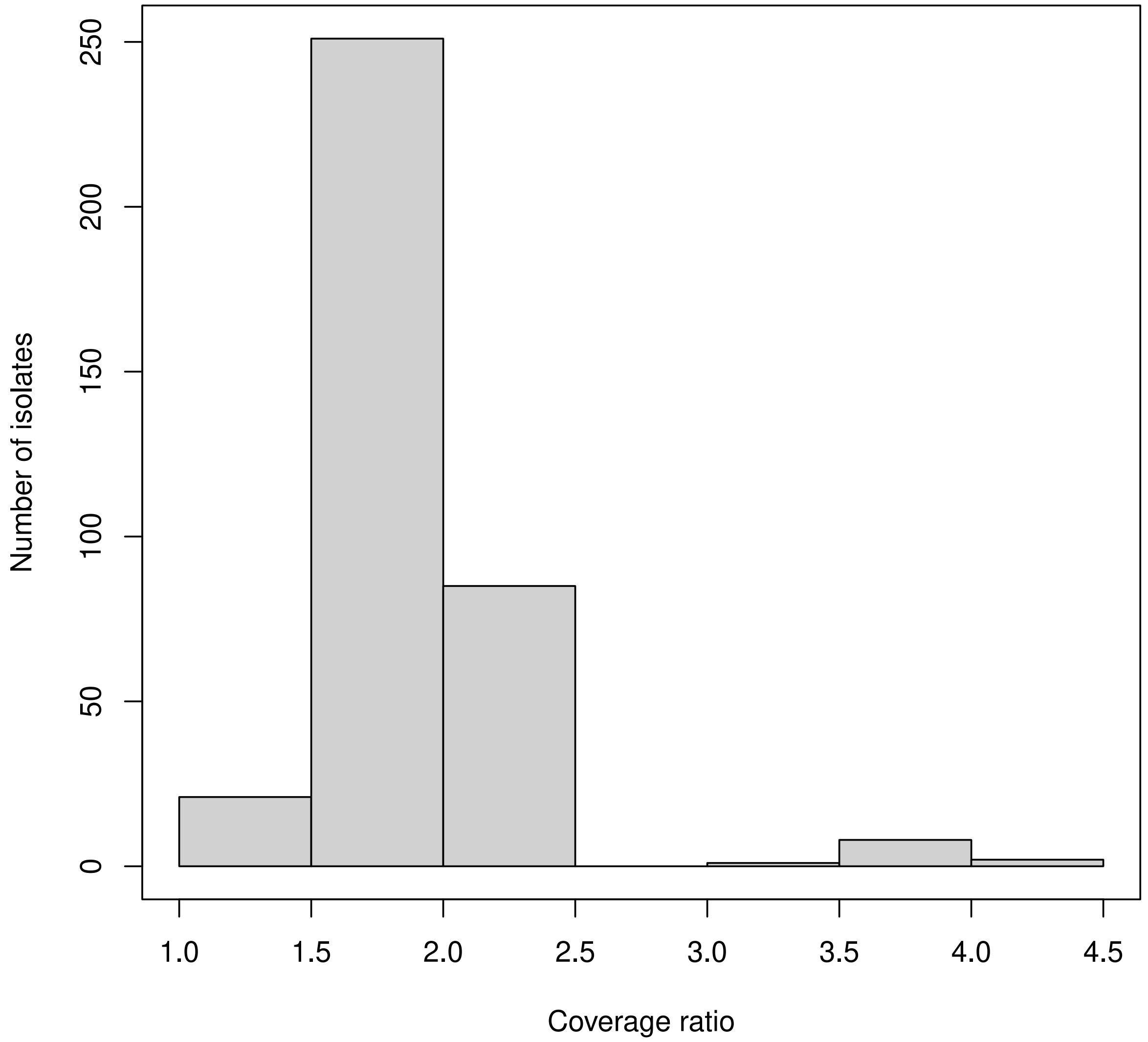

**S20 Fig. Copy number variation for the gene *AvrPm17***
Histogram of the ratio between the coverage of *AvrPm17* and the average genome wide coverage for all samples in the *Europe+_recent* dataset. A ratio between 0.5 and 1.5 is interpreted as one copy, between 1.5 and 2.5 two copies, and so on.

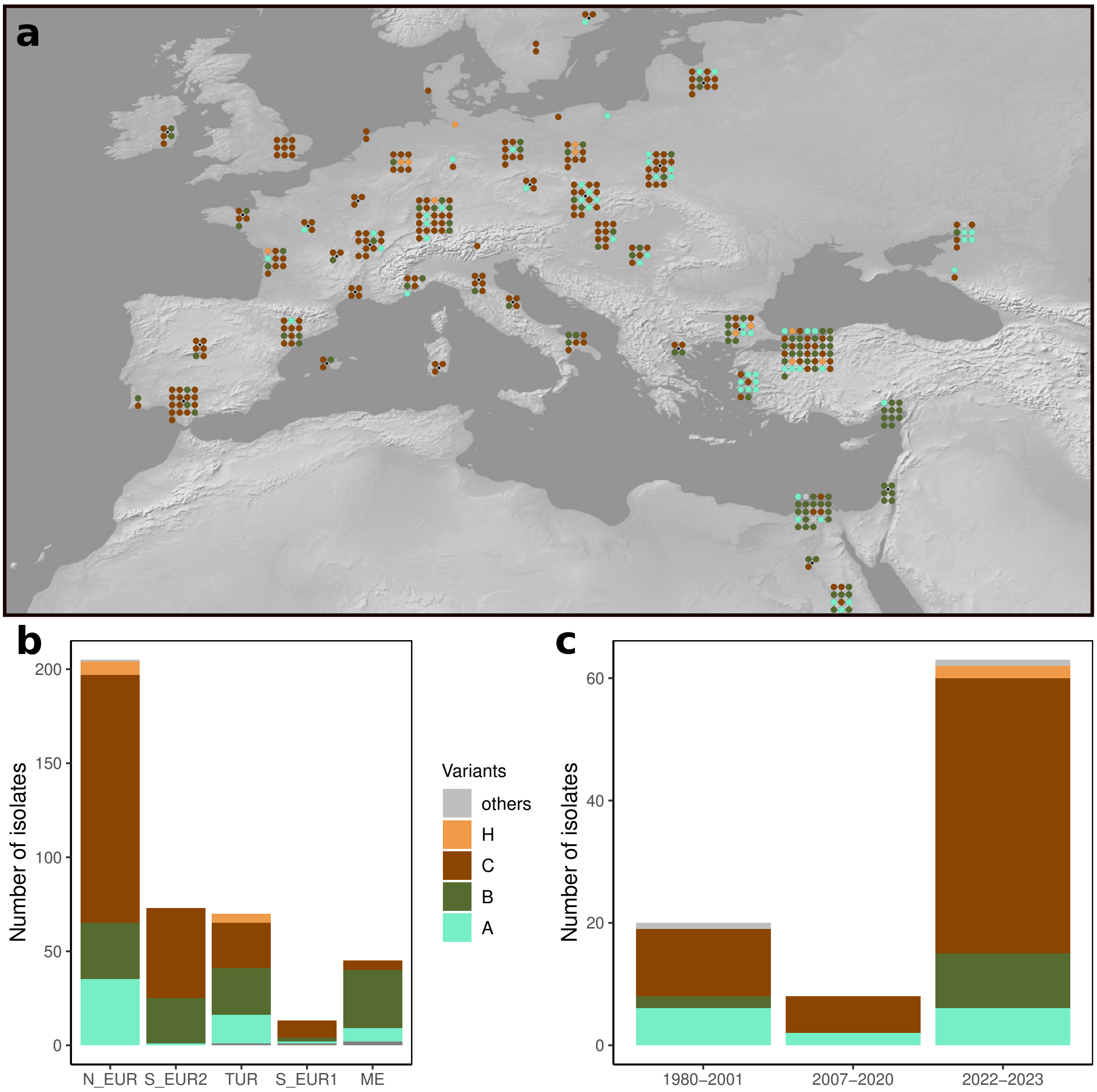

**S21 Fig. Distribution of AvrPm17 protein variants**
Protein variants are defined based on the mature protein sequence (after cleavage of the signal peptide). Same colours in all panels. If one isolate contains two different variants, both are depicted. (**a**) Map showing the distribution of variants in the *Europe+_recent* dataset. (**b**) Barplot showing the distribution of AvrPm17 variants in different populations (*Europe+_recent* dataset). (**c**) Barplot showing the distribution of variants in different time periods. For this analysis we focused on isolates sampled in Switzerland, the UK and France, as these were the only countries for which we had enough samples from different time points.

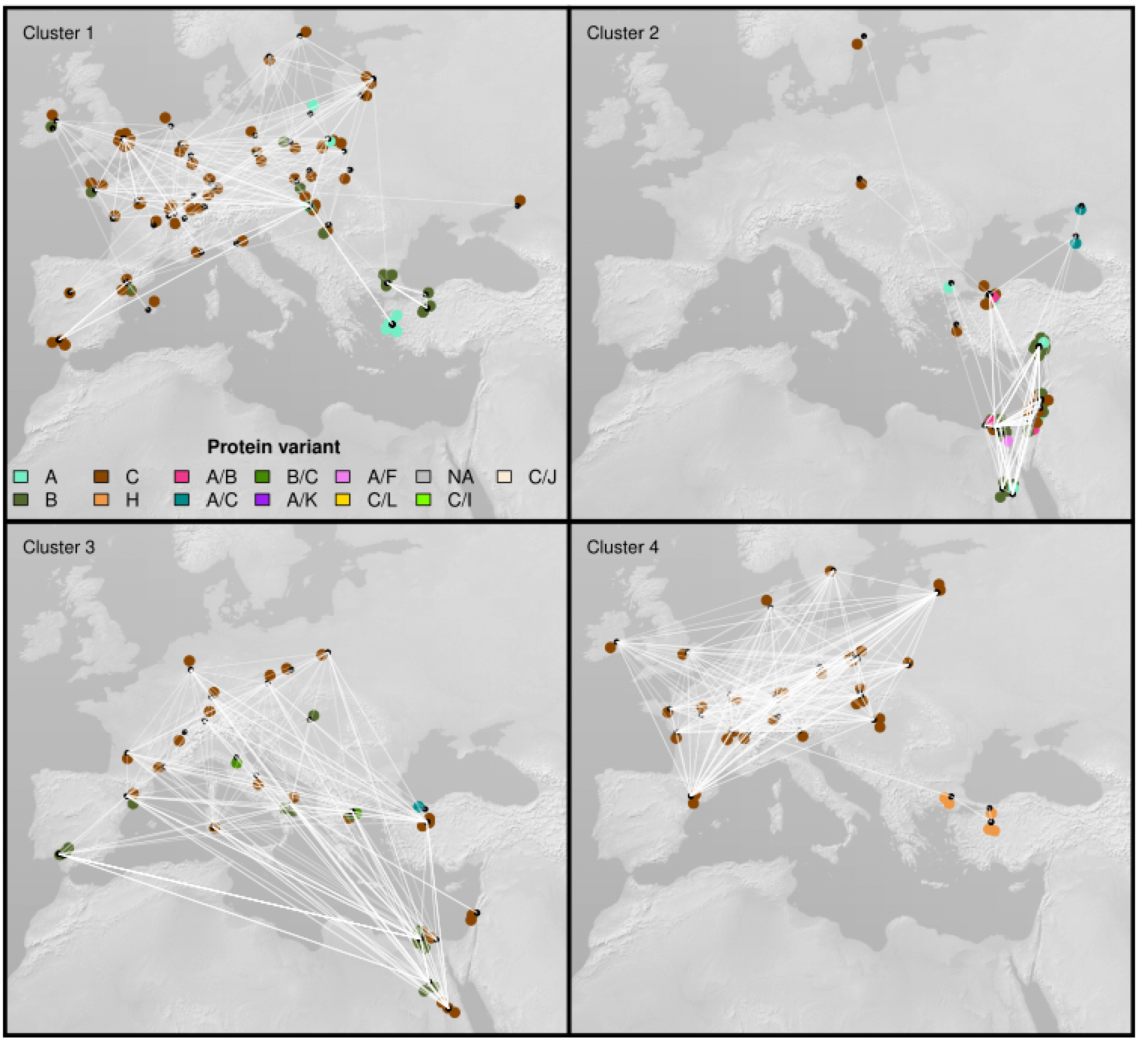

**S22 Fig. Geographic distribution of *AvrPm17* clusters 1-4**
The four largest clusters in the *AvrPm17* relatedness network are depicted separately. Small black dots represent sampling locations. Larger dots represent samples, and they are slightly moved from their sampling location to increase readability. Each sample is coloured based on the protein variant(s) coded by their respective *AvrPm17* genes. White edges connect isolates that are identical-by-descent over the *AvrPm17* locus.

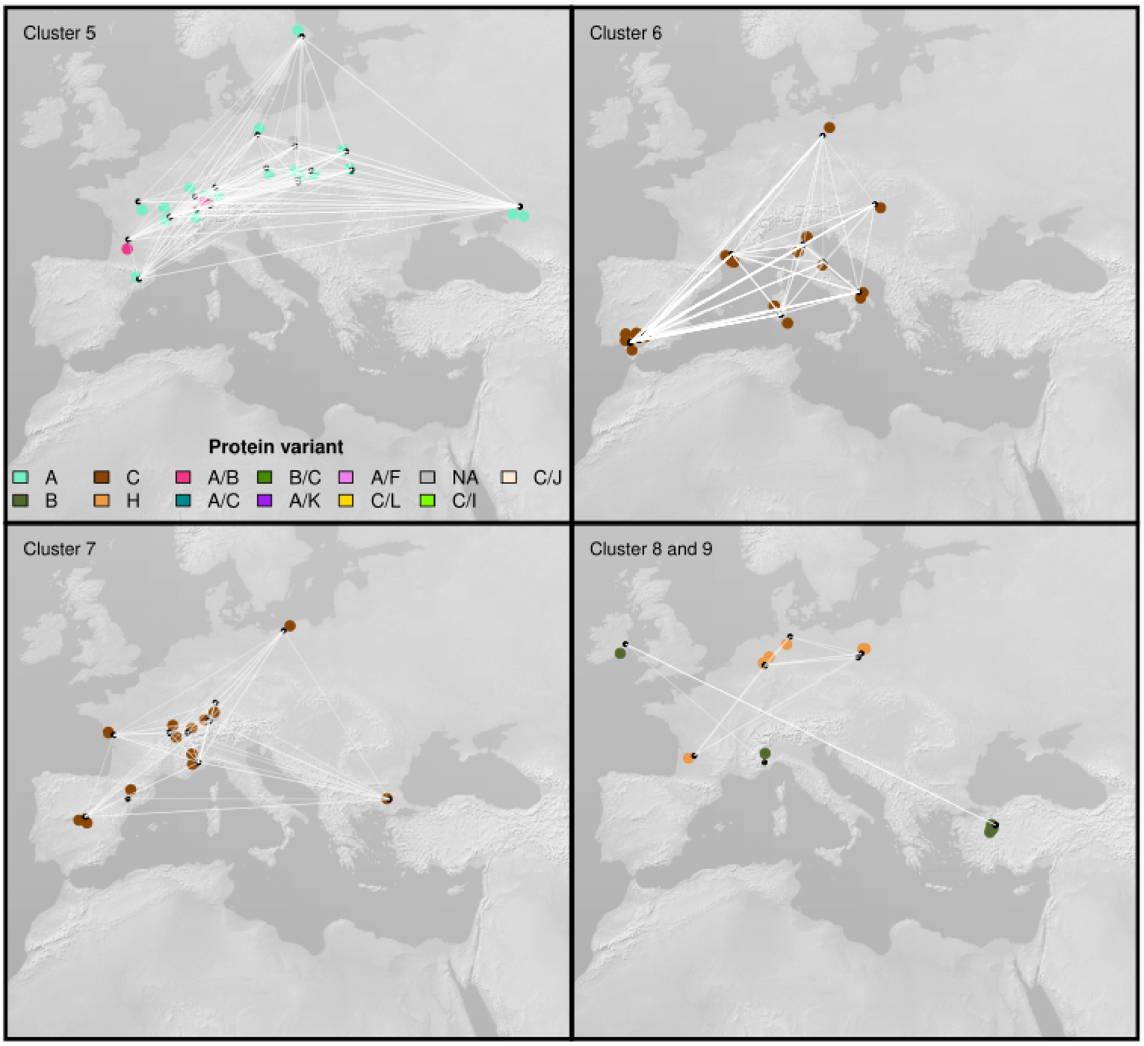

**S23 Fig. Geographic distribution of *AvrPm17* clusters 5-9**
The 5th, 6th and 7th largest clusters in the *AvrPm17* relatedness network are depicted separately. The 8th and 9th largest clusters are plotted in the same panel. Small black dots represent sampling locations. Larger dots represent samples, and they are slightly moved from their sampling location to increase readability. Each sample is coloured based on the protein variant(s) coded by their respective *AvrPm17* genes. White edges connect isolates that are identical-by-descent over the *AvrPm17* locus.

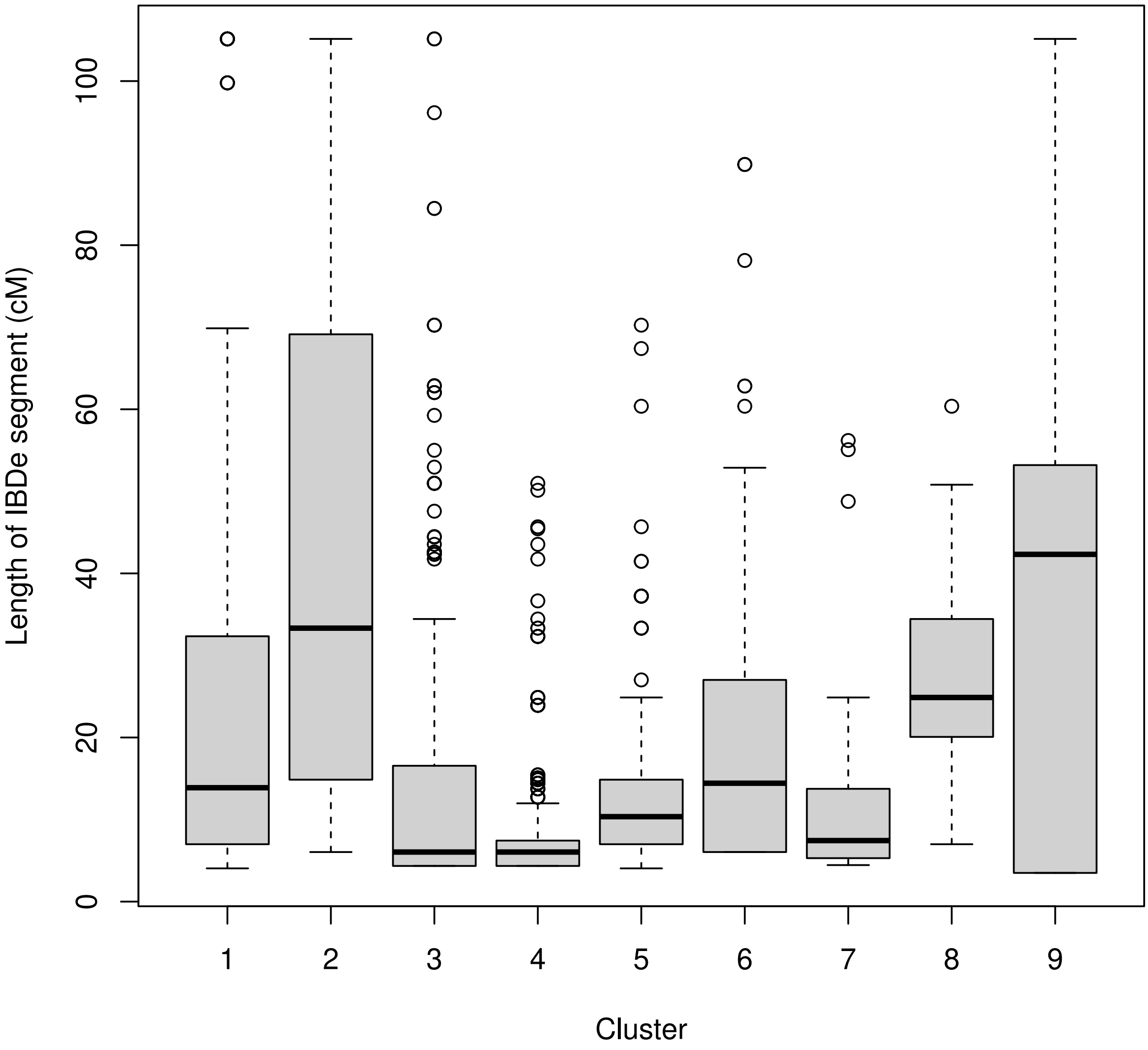

**S24 Fig. Distribution of lengths of identical-by-descent segments for the nine largest clusters**
The boxplots represent the distribution of the length of identical-by-descent (IBDe) segments in centiMorgans (cM) within each cluster. Larger segments correspond to more recent common ancestors. The average length in cM of an IBDe segment after x generations can be obtained with 100/2x(see **Methods**).

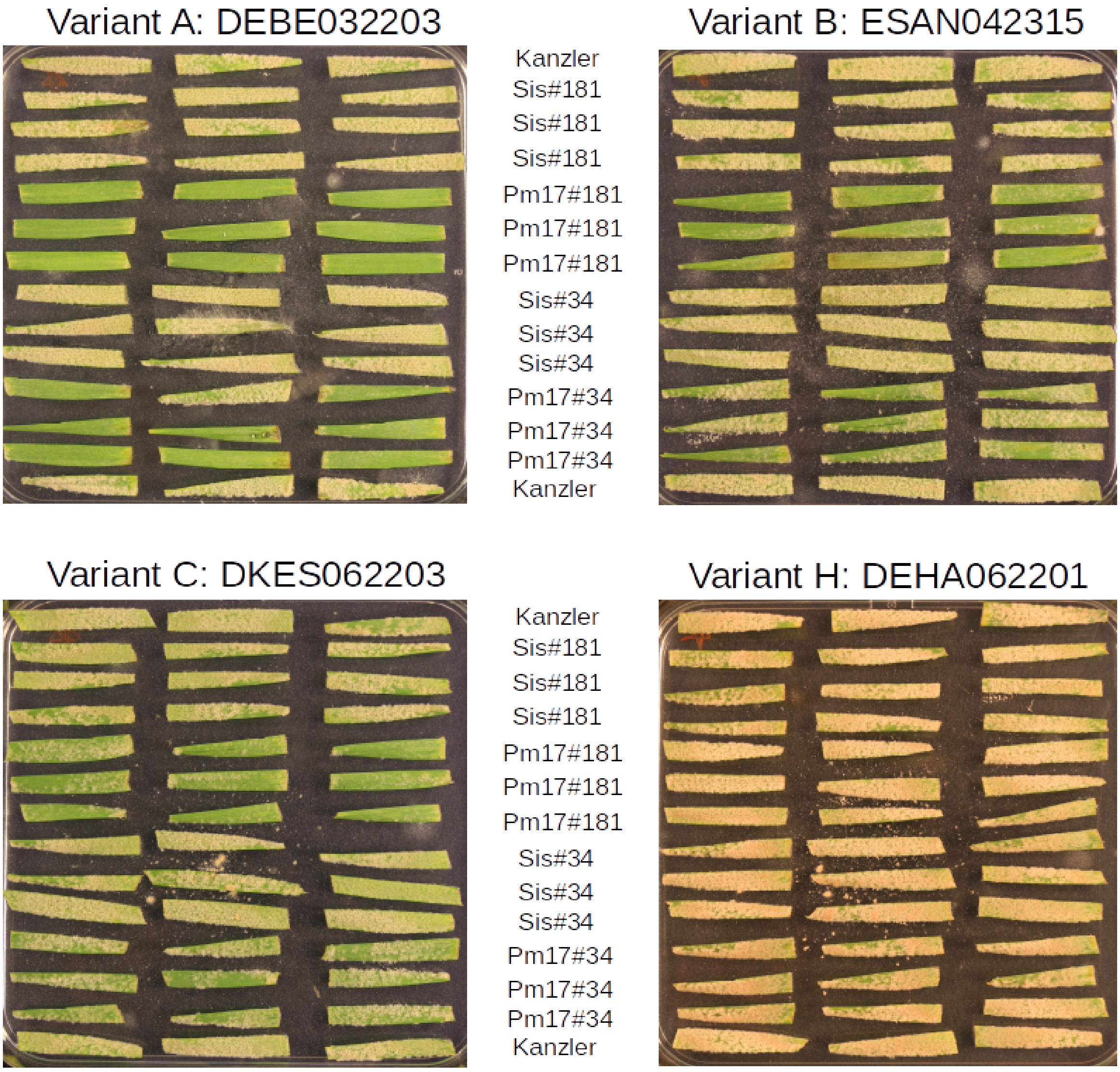

**S25 Fig. Infection tests on *Pm17* transgenic lines**

Eight Bgt isolates (four are depicted in **S25 Fig** and four in **S26 Fig**) from the 2022-2023 collection carrying different *AvrPm17* variants were tested for virulence on two transgenic wheat lines carrying *Pm17* (Pm17#181 and Pm17#34; see **Methods**). The corresponding sister lines (Sis#181 and Sis#34) and Kanzler, a susceptible cultivar, were included as controls. Each leaf fragment corresponds to a biological replicate. The isolate containing variant A is avirulent on *Pm17* lines, variants B and C are partially virulent, and variant H is completely virulent.

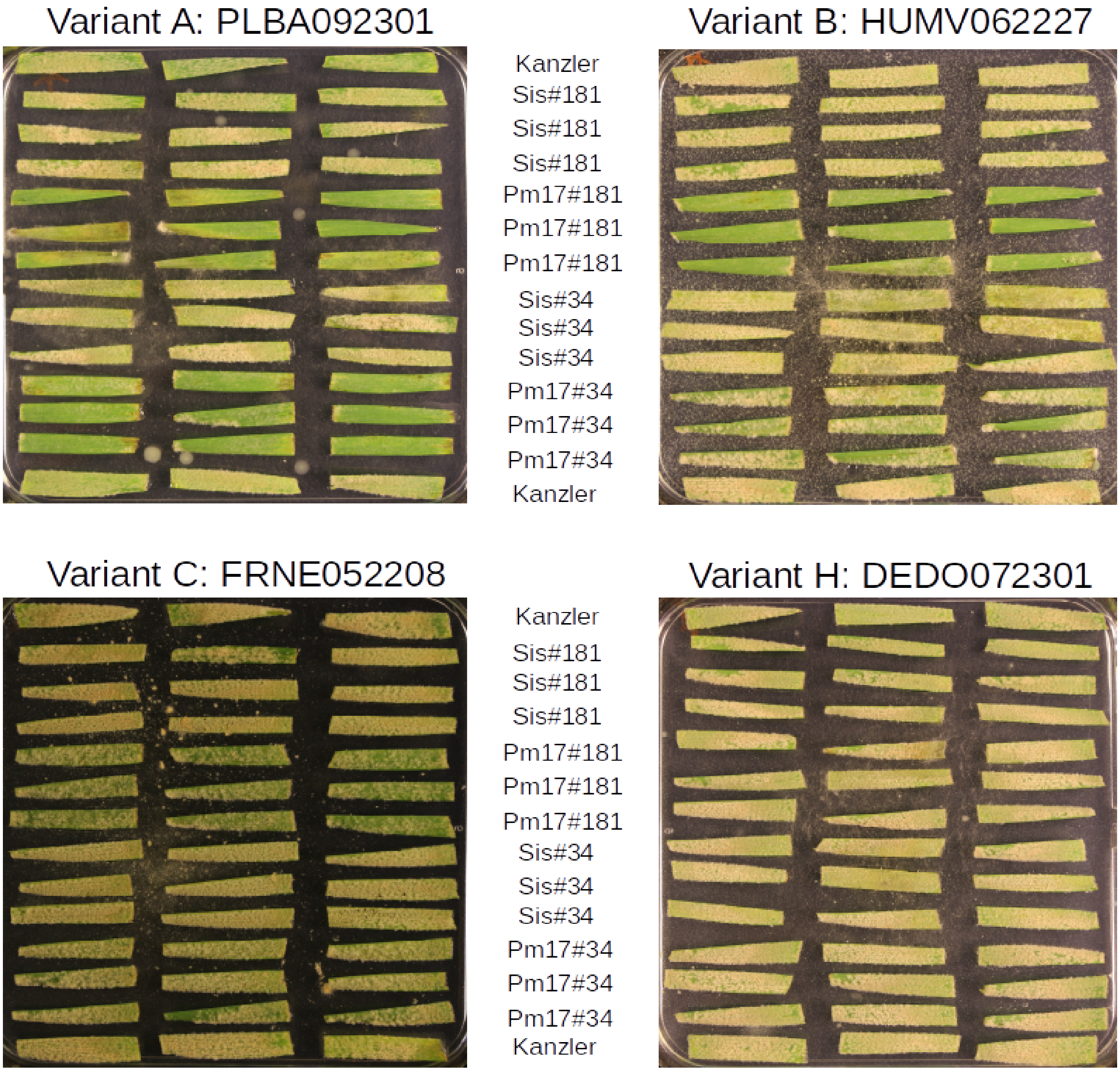

**S26 Fig. Infection tests on *Pm17* transgenic lines**
Eight Bgt isolates (four are depicted in **S25 Fig** and four in **S26 Fig**) from the 2022-2023 collection carrying different *AvrPm17* variants were tested for virulence on two transgenic wheat lines carrying *Pm17* (Pm17#181 and Pm17#34; see **Methods**). The corresponding sister lines (Sis#181 and Sis#34) and Kanzler, a susceptible cultivar, were included as controls. Each leaf fragment corresponds to a biological replicate. The isolate containing variant A is avirulent on *Pm17* lines, variants B and C are partially virulent, and variant H is completely virulent.

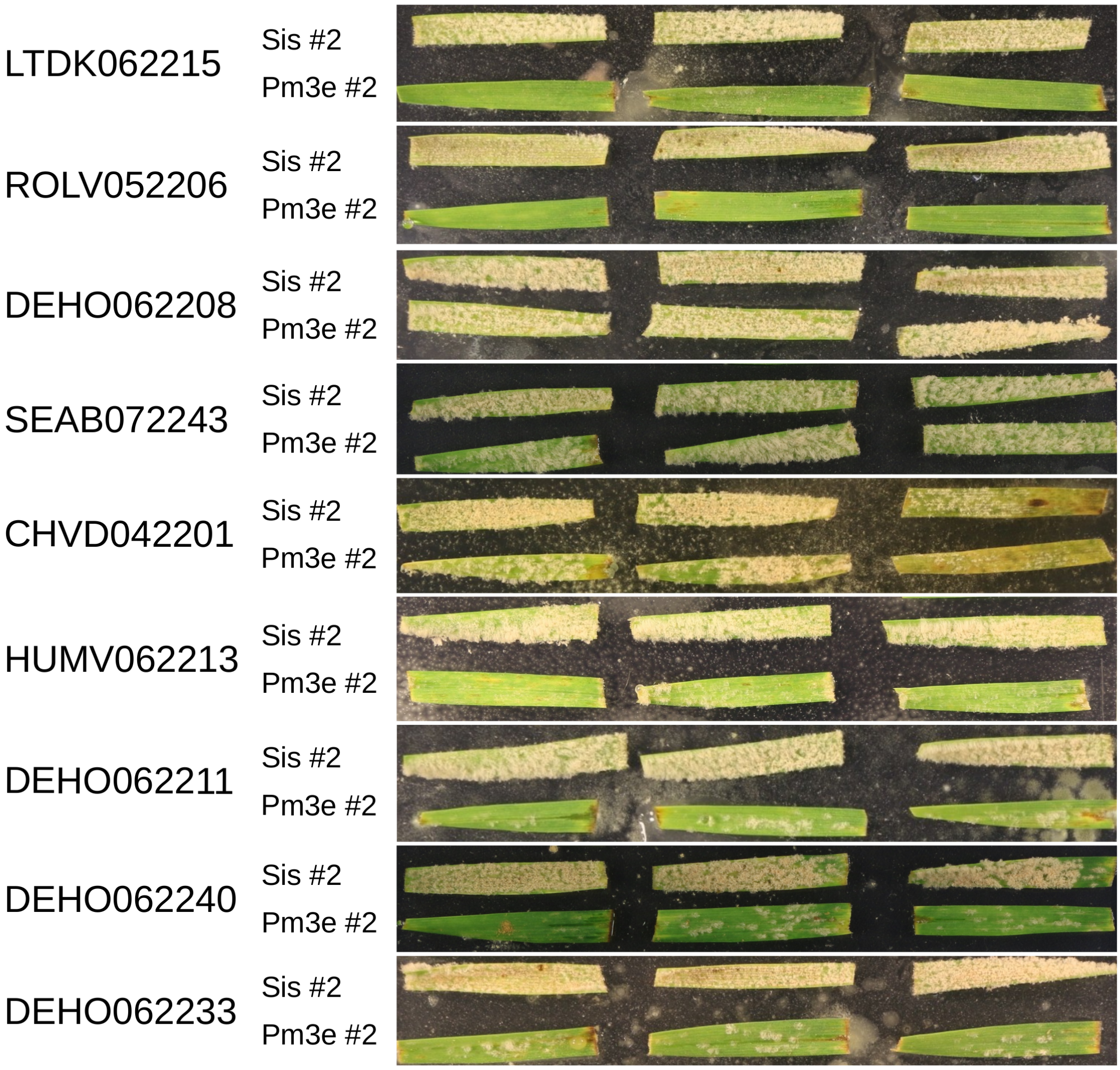

**S27 Fig. Infection tests on a *Pm3e* transgenic line**
Pm3e #2 is the transgenic wheat line containing the *Pm3e* gene and Sis #2 is the corresponding sister line used as a control. Each leaf fragment is a biological replicate. The first two isolates from the top (LTDK062215 and ROLV052206) show no signs of virulence on *Pm3e*. DEHO062208, SEAB072243, and CHVD042201 are fully virulent on *Pm3e*, while the remaining isolates are partially virulent.

### Appendices

#### S1 Appendix. Population structure analyses

##### Global population structure (*World* dataset)

One of our main objectives is to characterize the fine-scale population structure of wheat powdery mildew in Europe. However, as a first step we performed a population structure analysis of all available Bgt strains, independently of their time and place of sampling. Our goal with this analysis was to place the contemporary Bgt samples in the broader global context, and to identify eventual major changes in the global population structure. Such shifts are not unheard of. For example, previous population genetic studies of wheat yellow (stripe) rust (*Puccinia striiformis f. sp. tritici*) revealed that the European population was replaced by genotypes originating in eastern Asia, and that this occurred within a few years around 2011 [6, 7].

For wheat powdery mildew (*Blumeria graminis f. sp. tritici*, hereafter Bgt), it was shown previously that isolates sampled in different continents formed distinct populations [8]. More specifically, Sotiropoulos and colleagues showed that Bgt samples from Europe and the Middle East, which were mostly collected in the 1990s and the 2000s, grouped in two distinct clusters, clearly separated from isolates sampled in other regions (US, Argentina, China, Japan, and Australia; Figure 1 in [8]). In addition to the Bgt isolates from Sotiropoulos et al. [8] (except a few excluded by our quality filters), our World dataset (568 isolates in total, see **Methods**) contains 155 samples from North America, the Middle East and central Asia, collected between 2013 and 2018 [9], along with 255 isolates sampled by us in 2022-2023, in Europe and the Middle East. With the analysis of the World dataset, we want to test whether the recent isolates from Europe and the Middle East cluster together with the older ones originating from the same regions, and therefore whether there was a major shift or replacement of these Bgt populations.

We first performed a PCA which helped reduce the dimensionality of our data. We used biallelic SNPs and excluded singletons and sites with missing data exceeding 10%. PC1 explained 6.06% of the total variation while PC2 and PC3 explained 4.54% and 3.28% respectively (**S2d Fig**). As reported previously [8], we found that individuals from the USA, Japan, Argentina, and China formed distinct clusters, and that samples from Australia were indistinguishable from the USA isolates (**S2a-S2c Fig**). We also observed that the newly collected European and Middle Eastern samples clustered with the older samples from the same locations.

Next, we used ADMIXTURE [10] to study population structure by estimating individual ancestries. We performed this analysis for the complete *World* dataset to get an unbiased overview of relationships between samples from Europe and the rest of the world. The model was run for 10 different number of ancestries (K=1-10) for 10 replicates each and the "best" number of ancestries for our data was inferred by comparing the cross-validation errors for different values of K (**S3 Fig**). The first major drop in CV-error was found at K=4, where the four ancestries separated samples from four geographic regions (**S4a Fig**) (i) Northern/Central Europe, (ii) Southern Europe, Central Asia, and the Middle East, (iii) China and (iv) USA. The next few values of K separated the Caucasus and the Middle East from Southern Europe. Cross-validation errors for most replicates increased significantly from K=7 to K=10. However, the lowest cross validation error was obtained for one analysis with K = 9 (**S3 Fig**), and we used the ancestry proportions from this replicate for all further analyses. Under the best ADMIXTURE run, the nine ancestries defined nine groups of samples originating from (i) Northern/Central Europe, (ii) Southern Europe, (iii) Northern Turkey, Caucasus and Central Asia, (iv) Southern Turkey and Israel, (v) Egypt, (vi) China, (vii) Japan, (viii) the US and Australia, and (ix) Argentina (**S4a Fig**).

Overall, this analysis confirmed that Bgt populations in different continents are clearly distinct, and that the global population structure has remained stable in the last two to three decades.

#### Population structure in Europe and the Mediterranean (*Europe+* dataset)

We used 415 isolates collected between 1980 and 2023 from Europe and neighboring regions (*Europe+* dataset; see **Methods**) to explore finer-scale population subdivisions within the continent. We performed a PCA and found that the first three principal components explained 4.97%, 2% and 1.73% of the variation respectively (**S5d Fig**). PC1 separated the Northern/Central European isolates from the rest. PC2 and PC3 differentiated Egypt, Southern Europe, Israel and Turkey from each other (**S5a-c Fig**). Though the observed "clusters" are not entirely discrete, the PCA showed evidence for population subdivision within Europe, with the Northern/Central parts of Europe forming a distinct group, and Southern Europe and the Middle East showing further differentiation. Results of the ADMIXTURE analysis presented above were consistent with these findings and identified five major ancestries in our region of interest, roughly corresponding to (i) Northern Europe, (ii) Southern Europe (iii) Northern Turkey and Caucasus (iv) Southern Turkey and Israel (v) Egypt (**S4 Fig**).

We also observed varying levels of admixture across Europe and the Middle East (**S4 Fig**). Several samples from Northern Spain and Southern France showed a mix of Northern and Southern European ancestries. Some isolates from Northern Italy, Southeastern Europe and Greece showed admixture between the Turkish/Caucasian ancestry and the Northern European ancestry. In Egypt, some of the samples showed admixture with the Southern European ancestry, and some Israeli samples showed admixed Israeli and Egyptian ancestries.

To get a better resolution of the population structure, we used the haplotype-based method fineSTRUCTURE [11]. We used the *Europe+* dataset and filtered out all missing data which resulted in 1,201,198 biallelic SNPs. We computed the coancestry matrix, which is a representation of the ancestral relationships between samples (**S6 Fig**) and used it to cluster samples into populations. This method identified 45 populations in our dataset.

Within each population, members are statistically indistinguishable based on the coancestry matrix [11]. However, this criterion can over-split populations, generating subdivisions that are statistically valid, but have little biological meaning. To address this problem, and to investigate different hierarchical levels of structure, we inferred a population dendrogram, by repeatedly merging the two populations with the highest probability for the resulting merged group [11]. This method supported the main outcomes of the previous analyses. We observed that most isolates from northern and central Europe formed a large homogenous group separated from all other isolates. The samples from Southern Europe, the Middle East and the Caucasus were also clustered in separate groups, but with many finer, more pronounced subdivisions.

These three analyses (PCA, ADMIXTURE, and fineSTRUCTURE) identified the same main patterns in the data. Samples from northern Europe clustered together in a PCA, shared the same ancestry in ADMIXTURE, and constituted a large homogenous group in fineSTRUCTURE (**S4-S6 Fig**). All these results suggest a high rate of gene flow within this region. Samples from southern Europe, especially Spain and Italy, were separated from the rest by all analyses. In addition, fineSTRUCTURE revealed that isolates from Italy, Southern Spain, and Northern Spain formed distinct populations (**S7 Fig**). Compared to the north of Europe, isolates in these local populations shared more ancestry between them (**S6 Fig**), suggesting that they have smaller population sizes. Individuals from Turkey, Israel, and Egypt formed different groups in all analyses, and were also subdivided into several sub-populations. It must be noted, however, that the sampling scheme in these regions was clustered, with many samples collected from a few locations. The fine-scale subdivisions might thus be a result of this bias.

All three analyses identified several distinct groups, but these were not entirely discrete. We observed multiple intermediate individuals in the PCA, which had mixed ancestry in the ADMIXTURE results, and shared ancestry with individuals outside of their population in fineSTRUCTURE's coancestry matrix. Some examples are individuals in the north of Spain, in Southern France and in the north of Italy (**S4-S7 Fig**). These patterns are normally interpreted as the results of admixture events. Some of these samples might very well be admixed, in the sense that they descend from two recent ancestors with different ancestries. However, we think that the concept of admixture is a poor fit for an organism dispersed by wind, and we prefer to think of these as samples with intermediate genomic characteristics. One common limitation of ADMIXTURE and fineSTRUCTURE is that they assume a discrete number of ancestries or populations and no variability within them. These assumptions are violated when there are gradients of diversity over space, such as those typical of isolation by distance. It has been shown before that methods such as ADMIXTURE can infer multiple ancestries and admixed individuals to account for this variability [12]. These limitations were addressed by using different spatial population genetics approaches, which are reported in the main text.

### S2 Appendix. Redundancy analysis

Redundancy analysis (RDA), a statistical method that combines ordination and regression, is a powerful tool to test relationships between multivariate response and predictor variables such as in the context of genetic and environmental data [13]. We used RDA to infer the role of different environmental factors in shaping patterns of genetic diversity in European Bgt populations (*Europe+_recent* dataset). We included 12 climatic variables (filtered for multicollinearity), wind coordinates, geographical coordinates and country of sampling as explanatory variables (see **Methods**). We accounted for sampling coordinates and country separately to disentangle the effects of geography from country-specific agricultural practices. The full model (**genotypes** ∼ **climate** + **wind** + **geography** + **country**) explained 20% of the variation in the genetic data (**S7 Table**). We found that a large proportion of the variance in the model was confounded and could not be assigned to any predictor conclusively (36.6%). This is unsurprising, given the high levels of correlation expected among the explanatory variables. Nevertheless, all factors independently affected genetic variation, with climate and country contributing the most. This analysis suggests that local climatic conditions, agricultural practices, as well as wind and geography can partially explain genetic variation in Europe.

For an obligatory biotroph like wheat powdery mildew, genetic diversity may also be affected by its host. Two major kinds of wheat are cultivated in Europe: hexaploid bread and spelt wheat, and tetraploid durum/pasta wheat. There are regional differences in the extent of their cultivation, with most durum wheat being produced in Southern Europe and the Mediterranean, and hexaploid wheat in the north [14, 15]. These differences can potentially explain genetic differentiation in Bgt populations. We tested for this explicitly using 131 isolates that had been sampled from infected fields of known host types in 2022 and 2023. The RDA included host and the previously described predictors as covariates. The full model (**genotypes** ∼ **climate** + **wind** + **geography** + **country** + **host**) accounted for 31% of the variation in the data (**S8 Table**), and host alone had a small yet statistically significant contribution. However, as with the previous RDA, most of the variance in this model was confounded.

The power of the host RDA could be improved by adopting a different sampling strategy where Bgt is sampled from neighboring fields of hexaploid and tetraploid wheat that experience otherwise identical environmental conditions. Our dataset of 131 isolates includes a few individuals sampled from such locations in Spain, Italy and Hungary. Though samples from the Spanish and Italian locations showed no obvious signs of differential host adaptation (samples in close geographic proximity were also genetically very similar, irrespective of the host), those from Hungary stood out. We sampled 11 isolates of Bgt from nearby tetraploid and hexaploid wheat fields near Martonvásár (Hungary) over two years (2022-2023). Nine of these were sampled on hexaploid wheat and were found to belong to the N_EUR population. The remaining two were sampled on tetraploid wheat, one in 2022 and the other in 2023. They showed higher similarity to the southern European individuals and were classified as S_EUR2 by fineSTRUCTURE. The occurrence of different populations on different hosts is unlikely to be due to chance (Fisher exact test $p$-value = 0.0182), suggesting that wheat powdery mildew may exhibit host specialisation in some regions that should be investigated systematically with ad hoc sampling schemes.

Overall, we find that environmental conditions (climate, wind, geography) as well as the host can shape the landscape of genetic diversity of wheat powdery mildew in Europe. However, given the highly correlated nature of the predictors in our data, the effect of single factors could be disentangled only partially.

### S3 Appendix. Recent evolution, population genetics, and molecular epidemiology of AvrPm17

#### The 1AL.1RS translocation

The resistance gene *Pm17* was introduced into the gene pool of hexaploid wheat towards the end of the last century through the translocation of the short arm of rye chromosome 1 (1AL.1RS). In Europe, wheat lines containing the 1AL.1RS translocation were deployed only after 2000. While they were initially resistant to powdery mildew, their resistance was broken within a few years [2]. To check the current effectiveness of 1AL.1RS in Europe, we tested the virulence of Bgt isolates collected in 2022-2023 on the wheat line Amigo, the wheat variety in which 1AL.1RS was introduced from rye [16]. We found that 73.4% of them were at least partially virulent (138 out of 188 tested isolates), confirming that resistance conferred by 1AL.1RS was largely overcome in Europe (**Methods**, **S1 Data**). Virulence on 1AL.1RS should not be interpreted directly as virulence on *Pm17*. A previous study [2] found that at least one additional powdery mildew resistance gene was introduced into wheat with the same translocation. Therefore, 73.3% represents the minimum proportion of Bgt isolates that overcame *Pm17* resistance. Some of the remaining 26.7% are likely not virulent on Amigo because of the resistance provided by other genes. The genetic architecture of powdery mildew resistance in the wheat line Amigo was investigated by Müller and colleagues [2], and we refer to that study for further details.

#### The evolution and epidemiology of the four most frequent AvrPm17 protein variants

It was reported previously that Pm17 confers resistance upon the recognition of a specific Bgt effector (AvrPm17). We used coverage information to check how many copies of *AvrPm17* were present in the genome of each isolate. We found that in the *Europe+_recent* dataset, 326 isolates (88.6%) harbored two copies, 21 (5.7%) had one copy, and 11 (3.0%) had three or four copies (**S20 Fig, S1 Data**). We also found that in the 11 samples with more than two copies the nucleotide coding sequence was identical for all of them. While in 24.8% of the samples with two copies, the two nucleotide sequences were different. We classified all AvrPm17 sequences in protein variants based on the amino acid sequence of the mature protein (without considering the first 25 amino acids belonging to the signal peptide), and we followed the nomenclature of Müller et al. [2].

In total we observed nine variants (**S9 Table**), but only four were found in more than one or two isolates (variants A, B, C, and H), and in this section, we focus on those. We describe their geographic distribution within the *Europe+_recent* dataset, their temporal trajectories, and the evidence regarding whether they provide a fitness advantage against Pm17.

- **Variant A.** We found variant A in 16.3% of samples, making it the third most frequent variant. It is extremely rare in southern Europe, and more abundant in the other populations (**Fig 5, S21 Fig**). We tested how the frequency of variant A changed over time. For this analysis we considered isolates from France, Switzerland, and the UK, as these were the only countries from which we had enough samples from different periods. We found that before the introduction of *Pm17*, in the period 1980-2001, the frequency of variant A was 30%, while in the 2022-2023 collection its frequency decreased to 9.5%. This decline in frequency was statistically significant (Fisher exact test $p$-value = 0.034, odd ratio = 3.99; **S10 Table; S21 Fig**). This suggests a fitness disadvantage of variant A and was consistent with the results of its functional characterization [2]: variant A triggers a strong immune response when recognized by Pm17. Furthermore, isolates carrying this variant are avirulent on transgenic lines expressing Pm17 at high levels [2]. Surprisingly, we found that variant A was dominant in one of the large clusters (cluster 5), which is spread throughout the north of Europe, from France to Northern Caucasus (**Fig 5d**). This contrasts with the information presented above. Isolates carrying variant A are the least virulent on *Pm17* wheat lines, yet in cluster 5 this haplotype was highly successful. There are at least three hypotheses that could explain this pattern. 1) There could be a locus under selection in the vicinity of *AvrPm17*, and cluster 5 could be due to the selective pressure acting on this nearby locus (genetic hitchhiking). 2) Variant A could have provided a relative advantage compared to other variants that we cannot observe anymore. It is possible that other variants would trigger a stronger immune response compared to variant A, and that these have been selected out by the deployment of *Pm17*. Some good candidates for this hypothesis would be variants J, K, and L, which are each found in one copy in a single sample, and they all lack the A53V mutation that makes variants B and C less strongly recognized (**Fig. 5**). 3) It is also possible that variant A in cluster 5 was not selected, and that those isolates are identical-by-descent (IBDe) at the *AvrPm17* locus by chance. This cannot be excluded, as a significant excess of IBDe was not found in the Northern European population (N_EUR), to which most isolates in cluster 5 belong.

- **Variant B.** Variant B was the second most abundant (30.9% of samples), and it is prevalent in the Middle East and in Turkey (**Fig 5**; **S21 Fig**). We did not find it to have changed in frequency over time, nor to be over-represented in isolates that clustered in the relatedness network (**S10-S11 Tables**). Nonetheless, variant B is recognized less strongly by Pm17 compared to variant A, and isolates with variant B are partially virulent on *Pm17* transgenic lines [2].

- **Variant C.** Variant C is the most frequent variant, and it was found in 60% of Bgt isolates in the *Europe+_recent* dataset. It is the dominant variant in Europe, while it is less abundant in Turkey, Israel and Egypt (**S21 Fig**). It increased in frequency from 55% in 1980-2001 to 71.4% in 2022-2023 (in France, Switzerland, and the UK), although this difference could be explained by random sampling (Fisher exact test $p$-value =0.138; **S10 Table; S21 Fig**). In the relatedness network however, variant C was strongly over-represented in strains belonging to large clusters (**Fig 5**, **S11 Table**), indicating that it likely provided an advantage in the last two decades. Indeed, similarly to variant B, variant C can partially escape recognition by Pm17 (compared to variant A), and isolates carrying it are partially virulent on *Pm17* transgenic lines [2].

- **Variant H.** We found 12 isolates carrying a novel variant named variant H. All of them were collected after 2017 in northern Europe or Turkey. Variant H differs from variant C by one single amino acid change (Y31H). Interestingly the same mutation also occurred in a variant B background (generating variant I). The possible parallel evolution of the same mutation in two different backgrounds suggests that it might have a fitness advantage. Supporting this, all but one isolates carrying variant H belonged to a cluster (cluster 4 and cluster 8). Cluster 8 is composed of six isolates carrying variant H. The analysis of the length of IBDe segments showed that this is a recent cluster, and it might represent the onset of the expansion of a novel variant with a relative fitness advantage (**S24 Fig**). To test this hypothesis, we performed infection tests using the same two transgenic wheat lines used previously for the functional characterization of AvrPm17 [2]- Pm17#181 and Pm17#34. We tested two strains carrying variant H and found that they were fully virulent on the two *Pm17* lines (**Fig 6a, S25-S26 Figs**). We also repeated the *Agrobacterium* infiltration assay in *Nicotiana benthamiana,* which has previously been used to functionally validate AvrPm17 [2], and found that it cannot be distinguished from the negative control (**Fig 6b**), indicating that variant H is not recognized by Pm17 in this assay (see **Methods**).
